# Long-read sequencing of single cell-derived melanoma subclones reveals divergent and parallel genomic and epigenomic evolutionary trajectories

**DOI:** 10.1101/2025.08.28.672865

**Authors:** Yuelin Liu, Anton Goretsky, Ayse G. Keskus, Salem Malikic, Tanveer Ahmad, E. Michael Gertz, Farid Rashidi Mehrabadi, Michael Kelly, Maria Hernandez, Charlie Seibert, Juan Manuel Caravaca, Kayla Kline, Yongmei Zhao, Ying Wu, Biraj Shrestha, Bao Tran, Arindam Ghosh, Xiwen Cui, Antonella Sassano, Laksh Malik, Breeana Baker, Cornelis Blauwendraat, Kimberley J. Billingsley, Eva Perez-Guijarro, Glenn Merlino, Erin K. Molloy, S. Cenk Sahinalp, Chi-Ping Day, Mikhail Kolmogorov

## Abstract

Tumor evolution is driven by various mutational processes, ranging from single-nucleotide variants (SNVs) to large structural variants (SVs) to dynamic shifts in DNA methylation. Current short-read sequencing methods struggle to accurately capture the full spectrum of these genomic and epigenomic alterations due to inherent technical limitations. To overcome that, here we introduce an approach for long-read sequencing of single-cell derived subclones, and use it to profile 23 subclones of a mouse melanoma cell line, characterized with distinct growth phenotypes and treatment responses. We develop a computational framework for harmonization and joint analysis of different variant types in the evolutionary context. Uniquely, our framework enables detection of recurrent amplifications of putative driver genes, generated by independent SVs across different lineages, suggesting parallel evolution. In addition, our approach revealed gradual and lineage-specific methylation changes associated with aggressive clonal phenotypes. We also show our set of phylogeny-constrained variant calls along with openly released sequencing data can be a valuable resource for the development of new computational methods.

## Introduction

Tumor heterogeneity and evolution are driven by diverse genetic and epigenetic alterations, including single-nucleotide variants (SNVs), structural variants (SVs), copy number alterations (CNAs), and DNA methylation changes [106]. Traditionally, tumor evolution has been conceptualized as the gradual accumulation of point mutations under selective pressures [77]. More recently, distinct mutational processes that lead to the rapid accumulation of structural variants during catastrophic events — such as chromothripsis and chromo- plexy — have been increasingly recognized as critical drivers of tumor initiation and adaptation [96, 59]. Tumor evolution can also be shaped by non-genetic, or even non-heritable, determinants—especially epigenetic alterations such as DNA methylation or histone acetylation [100]. Profiling all these processes through a holistic approach is essential for improving our understanding of tumor evolution and for designing new treatment strategies [14, 5].

Short-read genome sequencing (e.g., Illumina) has been a workhorse of cancer genomics and has advanced our understanding of cancer heterogeneity and evolution [17]. However, these methods struggle to accurately detect structural rearrangements due to limitations in read mappability [48] or genomic coverage [109], and they require error-prone bisulfite conversion to detect epigenetic alterations [78]. As a result, short-read methods provide an incomplete view of the underlying mechanisms of tumor evolution.

Emerging long-read sequencing technologies, such as Oxford Nanopore Technologies (ONT) and PacBio, overcome the limitations of short-read sequencing by offering improved read mappability, direct variant phasing, and simultaneous capture of DNA methylation [58]. Several recent studies have used long-read sequencing to study complex patterns of somatic variation and methylation in various cancers [48, 24, 81], but so far mostly at bulk resolution.

To study tumor evolution on sub-clonal level, single-cell genome sequencing has recently been used to profile variants in individual tumor cells, however the technology is limited by sampling sparsity [26, 109]. Alternatively, whole genome sequencing of single-cell derived subclones have been used to precisely trace subclonal differentiation with high sequencing depth [102, 74, 13]. So far all such studies were based on short-read results, thus providing only partial view of genomic and epigenomic evolutionary landscape.

To address this issue, here we introduce an approach based on long-read sequencing of single-cell derived subclones; and apply it to analyze 23 melanoma subclones, each derived from a single cell of the B2905 cell line [84]—a murine model of UV-induced, RAS-mutated, and differentiation-transitory subtype of human melanoma. These subclones were previously characterized by differential therapeutic responses [34]; however, the contributions of SV and methylation changes to the genomic and phenotypic heterogeneity of these subclones remained unclear. We therefore generated high-quality SNV, SV, CNA, and differentially methylated region (DMR) calls in each subclone and developed new algorithms to infer their relative timing in distinct lineages through phylogenetic analysis.

Our long-read approach revealed the complex and multifaceted evolution of this heterogeneous tumor. We profiled the dynamic involvement of distinct mutational processes, including UV-induced damage, oxidative stress, and impaired DNA repair, offering deeper insight into their roles across the evolutionary stages of melanoma progression. We also observed recurrent amplifications of oncogenes occurring independently across multiple lineages and driven by distinct rearrangements, suggesting parallel adaptation under evolutionary pressures. Furthermore, we identified lineage-specific methylation trajectories associated with increased aggressiveness in distinct lineages, linking epigenetic changes with gradual phenotypic differentiation.

In summary, our approach that combines long-read sequencing with new and existing computational tools provides a highly detailed and holistic view of the evolutionary processes of a melanoma tumor. We also openly release the sequencing data and curated variant calls to encourage development and benchmarking of new computational methods.

## Results

### Long-read sequencing of single-cell derived subclones of B2905 mouse melanoma model

To showcase our long-read approach to profile tumor evolution at subclonal level, we used B2905 cell line, derived from M4 cutaneous melanoma model, previously characterized with high intratumoral heterogeneity and in preclinical studies showed a diverse response to immune checkpoint blockade (ICB) therapy [84]. Here we performed long-read sequencing of 23 single-cell derived subclones of the B2905 cell line, previously isolated for expansion to become individual clonal sublines, and implanted into C57BL/6 mice to observe distinct survival times [34]. While previous studies also analyzed bulk whole exome (bWES), whole transcriptome, and single-cell RNA sequencing (scRNA-seq) for each of the B2905 sublines [72, 40, 71], the contributions of SVs, CNAs and methylation to the evolution of this model remained unexplored.

We sequenced each subline using a single Oxford Nanopore Technologies (ONT) R10.4 PromethION flow cell. On average, this yielded 69.77Gbp of data per subline with a mean read N50 of 29.1Kbp and an average of 82.95% of base pairs with a quality score of 20+ (Supplementary Table S1). As a matching normal sample, we sequenced the splenocytes of a HGF-transgenic C57BL/6N mouse, the genetic background that was used to derive the B2905 model.

Following basecallling, we aligned the reads from each subline against the GRCm38 reference genome and to generate somatic SNVs, we used DeepVariant [86], and subtracted germline variants from the normal sample (Methods). For SV calls, we used Severus [48] in multi-sample mode (Methods). Additionally, since the mouse strain that derived B2905 cell line was generated by out-crossing HGF-transgenic FVB mice with C57BL/6 mice for over 10 generations, for SNV and SV variants, we subtracted known germline variants of the FVB mouse strain (Methods). Finally, we used Wakhan [2] for CNA calling, which uses SV breakpoint as the boundaries of CNA segments and fits a model integer copy number profile (Figure 1). Additionally, 5mC base modifications were profiled directly from the sequencing data, enabling the detection of DMRs using BSmooth [37], which we will discuss in further detail in a subsequent section. Together, these identified variants serve as the basis of the multifaceted analyses we will describe in the following sections.

**Figure 1:**
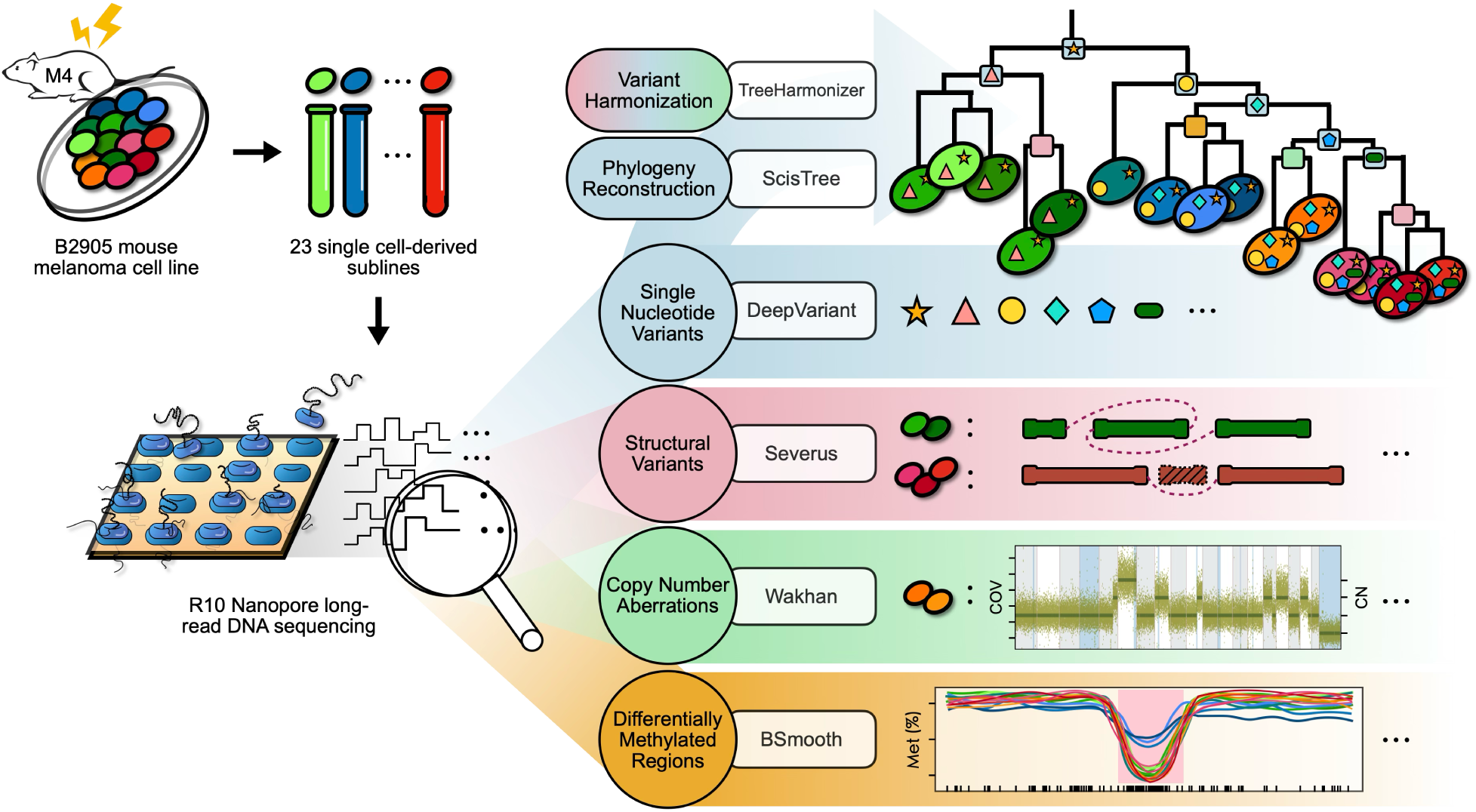
An overview of the long-read approach and analysis of 23 subclones of a mouse melanoma tumor. A tumor subline was cultured from each of the 23 single cells extracted from the B2905 cell line derived from the M4 mouse model of UV-induced melanoma, and the bulk samples were subjected to Nanopore long read sequencing. We studied melanoma tumor evolution integrating insights from mutation, structural variation, copy number variation, and differential methylation profiled from the sequencing data of the 23 sublines. Specifically, a tumor phylogeny of the sublines was constructed using SNVs, and then other types genomic and epigenomic events detected by various tools were placed onto the phylogeny. This provides an unprecedented look into melanoma evolution across time and various types of (epi-)genomic alteration events.

### Constructed phylogeny reveals phenotypically-distinct major clades

To infer the evolutionary relationship between subclones, we constructed a phylogenetic tree with ScisTree [72, 104] using a subset of high-confidence SNVs (Methods). From the initial set of 277,818 phylogenetically informative SNVs (61.75% of all SNVs used; excludes SNVs present in one or all sublines), we removed SNVs with low coverage (7.09% of informative SNVs) and SNVs that overlapped with deletions in any subline (89.00% of informative SNVs) (Methods). This resulted in a final set of 10,862 high-confidence, informative SNVs. The constructed tree consisted of 4 clades that we will refer as “major clades” - colored in green, orange, blue, and red - representing 4 distinct subclonal populations in the B2905 cell line (Figure 2a.).

**Figure 2:**
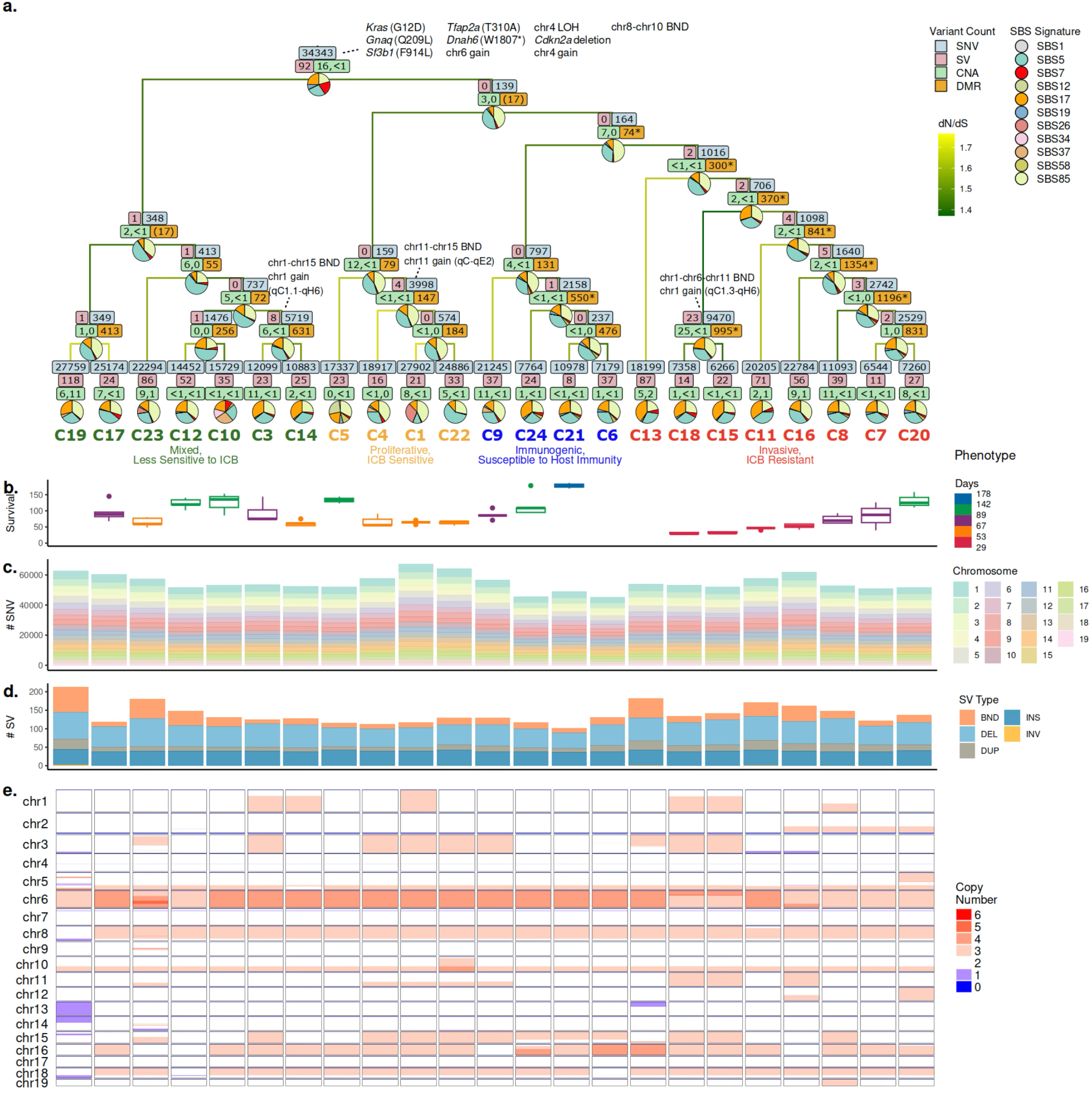
Phylogenetic tree and mutational landscape of 23 single-cell derived sublines. **a.** Phylogenetic tree with each node labeled with (i) counts of SNV, SV, CNA, and DMR assigned to the incoming branch of the node, (ii) exposure proportion of mutational signatures active along the incoming branch of the node, and (iii) key variants and events of note placed along the incoming branch of the node, if any. SNV and SV counts reflect those originating at that node. CNA numbers represent rounded percentages of genome collectively amplified and lost, respectively, among the subline below the branch, excluding those already accounted for in ancestral branches. DMR counts reflect the number of regions detected between the two groups of sublines defined by the bipartition of the tree at the node: the parenthesized numbers at the two immediate children of the root node reflect the same set of regions, and the numbers marked with an asterisk * denote those that are statistically significant per permutation tests (see Methods). The sublines are organized into 4 major clades, denoted by the distinct colors of the leaf labels. Sublines with previously determined phenotypes [40] also marked. **b.** Growth kinetic of tumors derived from individual sublines [34]. **c.** Per subline total SNV count per autosome. **d.** Per subline total SV count per SV type. **e.** Per subline copy number profiles.

In a parallel study, we characterized the contrasting phenotypes of selected sublines from each clades according to their *in vivo* growth rates, expression of cell cycle genes, and response rates to anti-CTLA4 ICB treatment [34, 40]. Sublines with zero or low tumor take rate in immunocompetant C57BL/6 mice could grow tumors in immunocompromised nude mice, suggesting that immunogenicity impacts *in vivo* growth of these sublines. Red clade had an aggressive phenotype with rapid *in vivo* growth and lower response to ICB, resembling invasive melanoma subtype [42]; while orange sublines had slower *in vivo* growth and higher ICB response, resembling proliferative subtype. Green sublines showed mixed phenotypes of red and orange clades. Blue sublines had low tumor growth in immunocompetent C57BL/6 mice are referred as immunogenic [19]. (Methods, Supplementary Table S2) Overall, phylogenetic reconstruction reflected the observed phenotypic differences between sublines and enabled the analysis of variant timing and lineage phenotypes associations that we describe below.

### Phylogeny-guided harmonization with **TreeHarmonizer** provides evolutionary timing of genomic variants

While the tumor phylogeny described above accurately reflects evolutionary relationship between the sub- lines, it excludes a large portion of SNVs that potentially overlap with deletions or have low coverage, and it does not include SVs or CNAs. This is typical for popular tumor phylogeny reconstruction methods that rely on SNVs [88] (see more detailed discussion in Methods). Inclusion of different variant types is complicated by possible overlaps (e.g. SNVs within subclonal deletions) or erronous variant calls. We thus developed a tool called TreeHarmonizer that (i) places variants on the branches of a phylogenetic tree and (ii) identifies variants in disagreement with tree.

Briefly, TreeHarmonizer takes as input (i) a phylogenetic tree and (ii) individual SNV, SV, and CNA calls for every subline. For each unique variant, it evaluates whether the variant is consistent with the input phylogeny based on (i) the most-recent common ancestor (MRCA) of all sublines carrying the variant, (ii) the variant status in all sublines, and (iii) the presence of nearby deletions (Methods). We denote a variant as clonal, subclonal, or private, if they were placed at the trunk, internal branch, or terminal branch of the tree, respectively (Figure 2).

On our mouse melanoma dataset, TreeHarmonizer placed high proportions of all SNVs and SVs (96.71% and 95.49%, respectively). Of placed SNVs, 7.89%, 8.38% and 83.73% were clonal, subclonal and private, respectively. Within individual sublines, on average 62.23%, 8.69% and 29.08% SNVs were clonal, sub- clonal and private, respectively (Supplementary Table S3). Proportionally, SVs were distributed similarly to SNVs (Supplementary Table S4). The variant allele frequency (VAF) of placed clonal and subclonal variants was distributed around 50%, as expected for heterozygous somatic variants fixed in a given lineage (Supplementary Figure S2c.,d.). Overall, high proportion of placed variants confirms the agreement between the phylogenetic tree and genomic landscape of the sublines. It also suggests that sublines were free of contamination, as it would result in an increase in unplaced variants.

To further evaluate the correctness of variant placement, we examined the set of SNVs that were placed by TreeHarmonizer differently, compared to the initial phylogenetic assignment by ScisTree, which typically relies on more stringent filters for input variant calls in searching for the correct tree structure (Methods). We compared the partition function values [71] that estimates the probability that, given a set of calls in individual sublines, a mutation is placed at the MRCA branch of a clade (Methods). Indeed, TreeHarmonizer variant placement led to a significant increase (one-tailed paired-sample *t*-test t(26) = 11.99*, p* = 2.133*e* − 12) in their partition function values, confirming accurate variant placement (Figure 7g.).

### Evolutionary dynamics of somatic SNVs and mutational processes

Overall, the cumulative number of variants (all variants acquired along the course of evolution at a given tree node) was variable between sublines and major clades; in particular, sublines in the blue clade tend to have below-median number of both SNVs and SVs (Figure 2c.,d., Supplementary Figure S2a.). The number of SNVs and SVs within individual branches in the tree were strongly correlated (*r*(43) = .798, *p* = 5.158*e* − 11 *< .*001; Supplementary Figure S2b.), suggesting that the rate of SNV acquisition relative to that of SV is roughly constant over time.

Given the number of variants placed by TreeHarmonizer, we sought to explore the timing of mutational processes, commonly explained through the patterns of somatic SNVs. We estimated activities of COSMIC [93] mutational signatures at each branch of the tumor phylogeny (Figure 2a.; Methods). Notably, SBS7 (a UV damage signature) was prevalent at the trunk (Figure 2a., Supplementary Figure S8), consistent with the fact that the tumor model was induced by UV radiation.

Some signatures were present throughout the phylogeny, such as clock-like signatures (SBS1 and SBS5), and SBS85 (attributed to activation-induced cytidine deaminase; AID) (Figure 2a., Supplementary Figure S8). While clock-like signatures presence is expected, SBS85 has been typically described in the context of lymphoid cancers [47]. However, it has also been shown that inflammation-induced AID expression in non-lymphoid cells promotes skin cancer development independently of UV damage [76].

Other signatures were more specific with timing or lineages. For example, SBS26 (defective DNA mismatch repair), had increased activity solely at the terminal branch attached to subline C1, which had the highest cumulative number of SNVs among all sublines (Figure 2a.,c., Supplementary Figure S8). SBS17, a signature potentially associated with damages inflicted by reactive oxygen species (ROS), shows significant difference in activity both between internal and terminal branches (adjusted *p* = 9.08e-05; Methods), and between branches in the orange and red clade (adjusted *p* = 9.68e-03; Methods, Supplementary Figure S8). While the differences between internal and terminal branches could be explained by the additional oxidative stress caused by cell culture [36], the difference between the orange and red clade may reflect the difference in the metabolism between them, as we have shown previously that the expression of mitochondrial genes was selected in the former [40]. (Figure 2a., Supplementary Figure S8).

To identify putative initiating events and variants that contributed to lineage differentiation, we performed functional annotation and enrichment analysis of SNVs (Methods, Supplementary Figure S7). Interestingly, dN/dS values at internal branches were significantly lower than terminal branches (*p*=5.9e-07, Supplementary Figure S7a., Methods), suggesting an increasing positive selection pressure over the evolutionary timeline (Figure 2a., Methods). Among the clonal SNVs, we observe hotspot coding mutations, such as *Kras* (G12D) and *Gnaq* (Q209L) that are well-known cancer drivers, and variants in *Sf3b1* and *Tfap2a*, which are frequently mutated genes in melanoma [75, 45]. (Figure 2a.). Overall, impacted genes were enriched in pathways related to neural system development and differentiation as well as differentiation of neural crest to melanocytes (Supplementary Figure S7b.). At the same time, individual clades exhibited enrichment in functionally distinct and phenotypically relevant pathways, such as DNA damage checkpoint signaling in the green clade (Supplementary Figures S7c.), cell cycle and metabolism in the orange clade (Supplementary Figures S7d.) and cell migration and methylation in the red clade (Supplementary Figures S7f.). Overall, this analysis revealed the dynamics of mutational processes and putative functional effects of lineage-specific mutations, that we associate with other genomic, epigenomic or phenotypic changes below.

### Somatic SVs dynamics suggest early drivers and late genomic instability

Long-read sequencing enables accurate detection of somatic SVs, which may be driven by orthogonal mutational processes and have substantial impact on lineage phenotypes, as we illustrate below. Overall, small deletions (*<*1kb) were the most common SV; insertions were also common at the clonal level, but less common as private; and (interchromosomal) breakend junctions were more common at the private level (Figure 3a.-d.). Severus further groups breakend junctions into clusters if they are likely to result from a single mutational event. We identified 65 such clusters involving a total of 298 SVs: of which four were composed of clonal SVs 3e.), one of subclonal SVs (e.g. Figure 3f.), and 60 of private SVs (e.g. Figure 3g.). Private SV clusters were the most complex: while clonal clusters consisted only of two SVs each, 14 private clusters had at least five SVs, and largest cluster had 28 SVs (in C13). The green clade was overall enriched with clustered SVs, compared to the other clades (150/298 clutered SVs were in the green clade; Fisher’s exact test, p = 3.055e-07), primarily driven by C19 that had the highest number of clustered SVs (n = 68) in nine clusters (Supplementary Figure S3). Overall, the vast majority of clustered SVs were private, suggesting that in this tumor complex rearrangements were late events, potentially driven by an increased genomic instability.

**Figure 3:**
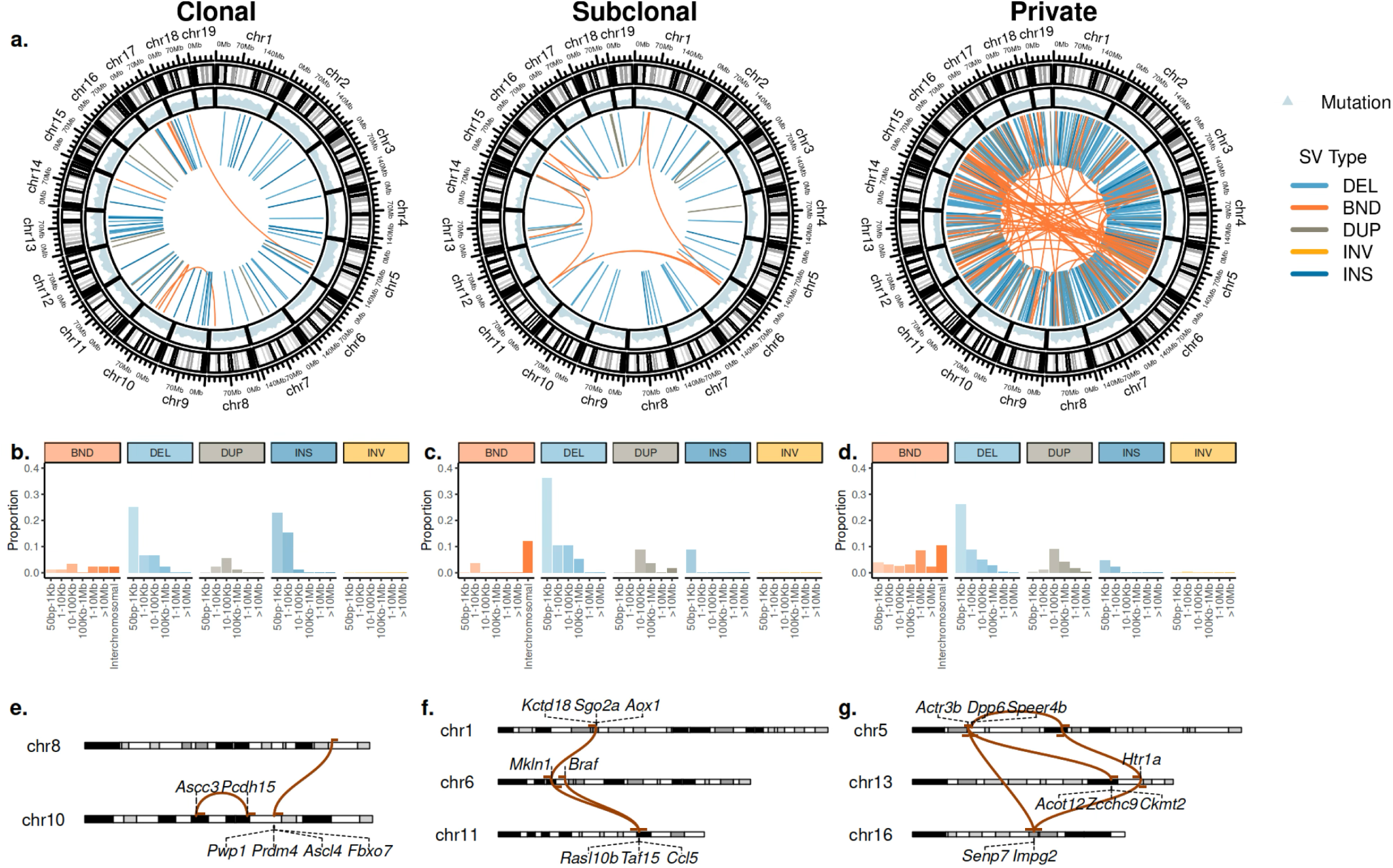
Patterns and dynamics of somatic SVs. **a.** Chromosomal distribution of clonal, subclonal, and private SNV and SV events. **b-d.** Distribution of clonal (b.), subclonal (c.), and private (d.) SVs across categories defined by SV type and size. **e-g.** Examples of notable clonal (e.), complex subclonal (f., shared by sublines C15 and C18), and complex private (g., private to C10) interchromosomal SVs. Genes near breakpoints are annotated.

Using Padfoot (https://github.com/KolmogorovLab/Padfoot), we identified 564 SVs with breakpoints occurring within gene boundaries (Methods). Of these, 330 affected coding regions through exon deletions or disruptions involving exonic sequences. Notably, we observed clonal deletions affecting exon 4-9 of *Nav3* (Supplementary Figure S9a.), which included the first coiled-coil domain, the mutations in which have been shown to result in a loss of function [32]. In accordance, deletion of *NAV3* gene has been linked to poor prognosis in cancer [6]. Additionally, in sublines C3 and C14, we identified a subclonal deletion in *Met* resulting in the loss of exon 4-11 (Supplementary Figure S9b.), which included extracellular and regulatory domains. Such truncated *MET* has been shown to be self-activated and promoting immortalization of normal cells [3]. We also detected 24 candidate gene fusions, of which 23 were private. Among them, a subclonal fusion between *Map4* and *Smarcc1* was identified in C3 and C14 and validated using scRNA-seq (Supplementary Figure S9c.). Beyond disrupting genes and generating oncogenic fusions, SVs also cause CNAs that alter gene expressions, which will be the focus of the following subsection.

### Recurrent CNAs in independent lineages suggest parallel adaptation process

SVs often cause CNAs potentially affecting large contigous regions of the genome. Overall, a high proportion of the genome was impacted by amplifications (26.37% on average per subline), and smaller fraction impacted by losses (1.24%) (Figure 2a.,e.). To understand functional implications of these alterations, we focused on genes with one-to-one human homologs curated in the COSMIC Gene Census [93] and labeled them as oncogene (ONC), tumor suppressor gene (TSG), or not designated (ND; Methods, Figure 4, Supplementary Figures S10-S16).

**Figure 4:**
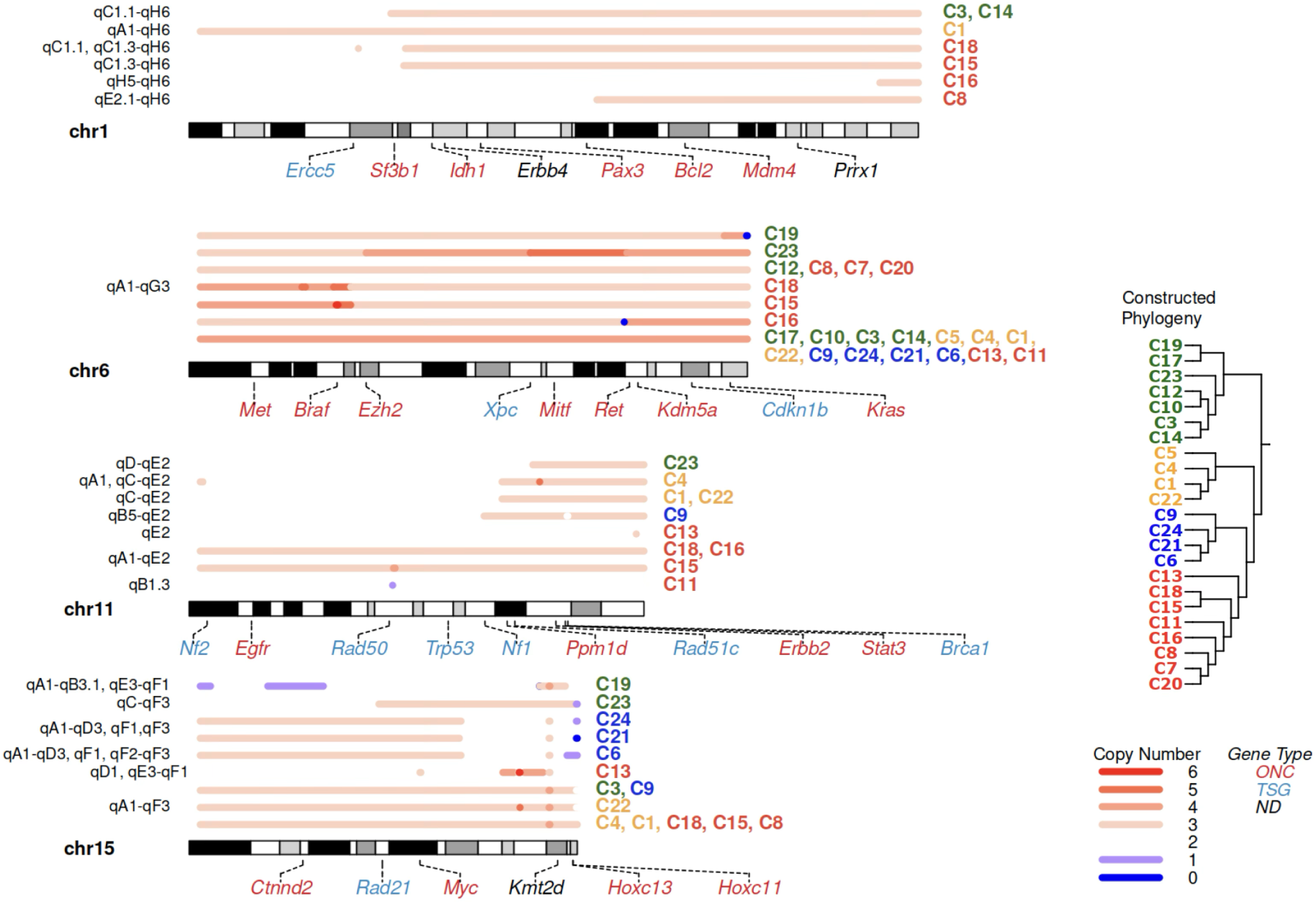
Genomic regions recurrently affected by CNAs suggest parallel evolution. Examples of regions on chr1, chr6, chr11, and chr15, recurrently harboring CNAs, possibly due to independent genomic events that impact the same region. Within each chromosome, sublines harboring CNAs are grouped by the similarity of their copy number profiles, and the respective copy number profiles are annotated by the chromosome bands they span. Select genes are annotated along the chromosomes and colored by whether their human homologs are designated as oncogene (ONC) or tumor suppressor (TSG) according to COSMIC Gene Census [93].

We first focused on clonal CNAs, such as whole-chromosome amplification of chr6 in all sublines, although some sublines acquired additional copies in later events (Figure 4). These CNAs affected key ONCs, such as *Met*, *Braf*, *Mitf*, and clonally mutated *Kras* (Figure 4, Supplementary Figure S12). We also observed a focal loss of of *Cdkn2a* in both copies of chr4 with the same breakpoint boundaries. This was likely a result of focal deletion, followed by copy-neutral loss of heterozygosity (LOH) in a larger region, happened approximately half way through the phylogeny trunk based on variant timing analysis (Methods). These overall suggests a substantial number of early CNA events could have contributed to tumor initiation.

Next, we analyzed the patterns of subclonal and private CNAs. While many of them were attributed to internal tree branches by TreeHarmonizer, others remained unplaced. In unplaced CNAs, similar regions were amplified by different SV breakpoints in independent parts of the tree (Methods). This suggests that these amplifications were independent recurrent events in different lineages, representing examples of parallel evolution. In one case, five distinct genomic rearrangement events resulted in independent amplifications of a large portion of chr1 in seven sublines (Figure 4): two SVs were subclonal, three were private; most of them were products of intra-chromosomal fusions, with one exception of whole-chromosome amplification in C1. In another example of parallel CNA, a chr11 gain in C4, C1, and C22 is associated with the same subclonal interchromosomal SV involving chr15, the one in C9 involves chr18 and results in the amplification of TSG *Nf1*, which also carries a private, non-synonymous SNV in C9 (Figure 4, Supplementary Figure S14). This analysis suggests that subclonal CNAs substantially contributed to differentiation of tumor clades, and indicates convergent evolution in independent lineages.

Indeed, among COSMIC genes impacted by CNAs, ONCs were preferentially gained (*p* = 3.832e-02, Methods) and TSGs were preferentially lost (*p* = 7.294e-03, Methods) in this tumor, as expected ; suggesting that those changes may be independently selected for in a highly heterogeneous tumor [50]. For example, parallel amplification of *Myc* was recurrently present in a majority of sublines (Figure 4, Supplementary Figure S15); likewise, *Myc* homolog is commonly amplified in human melanoma [95]. In C13, *Myc* is duplicated via a focal SV spanning merely 1.6Kbp, likely reflecting targeted selection. Furthermore, a closer examination of ONC/TSG COSMIC gene homolog pairs adjacent to copy number change boundaries shows that, more often than expected by chance, ONC falls on the side of the boundary with the higher copy number state while TSG on the lower side (*p*=3.74e-02, Methods). This provides evidence that relative positioning of ONC and TSG genes on chromosome may influence the selection of CNA boundaries.

### Methylation changes contribute to differentiation of an aggressive clonal lineage

In addition to genomic variants, epigenomic changes may shape the observed distinct lineage phenotypes. Long-read sequencing, combined with phylogenetic approach, enabled us to identify genomic regions that are differentially methylated at subclonal resolution (Figure 2a).

For each internal branch in the phylogeny, a set of DMRs between the bipartitions of the sublines induced by the corresponding tree split was identified using BSmooth [37, 29]. We then evaluated significance of the number of DMRs obtained via a permutation test for branches ancestral to a particular number of leaves (Methods, Supplementary Figure S18). We found a strong positive correlation between the number of cumulative SNVs and the number of DMRs at the branch (*r*(5) = .842*, p* = 1.758e-2, Methods; Figure 5a.), suggesting that the rate of differential methylation to some extent mirrors that of SNVs in the evolution of this tumor.

**Figure 5:**
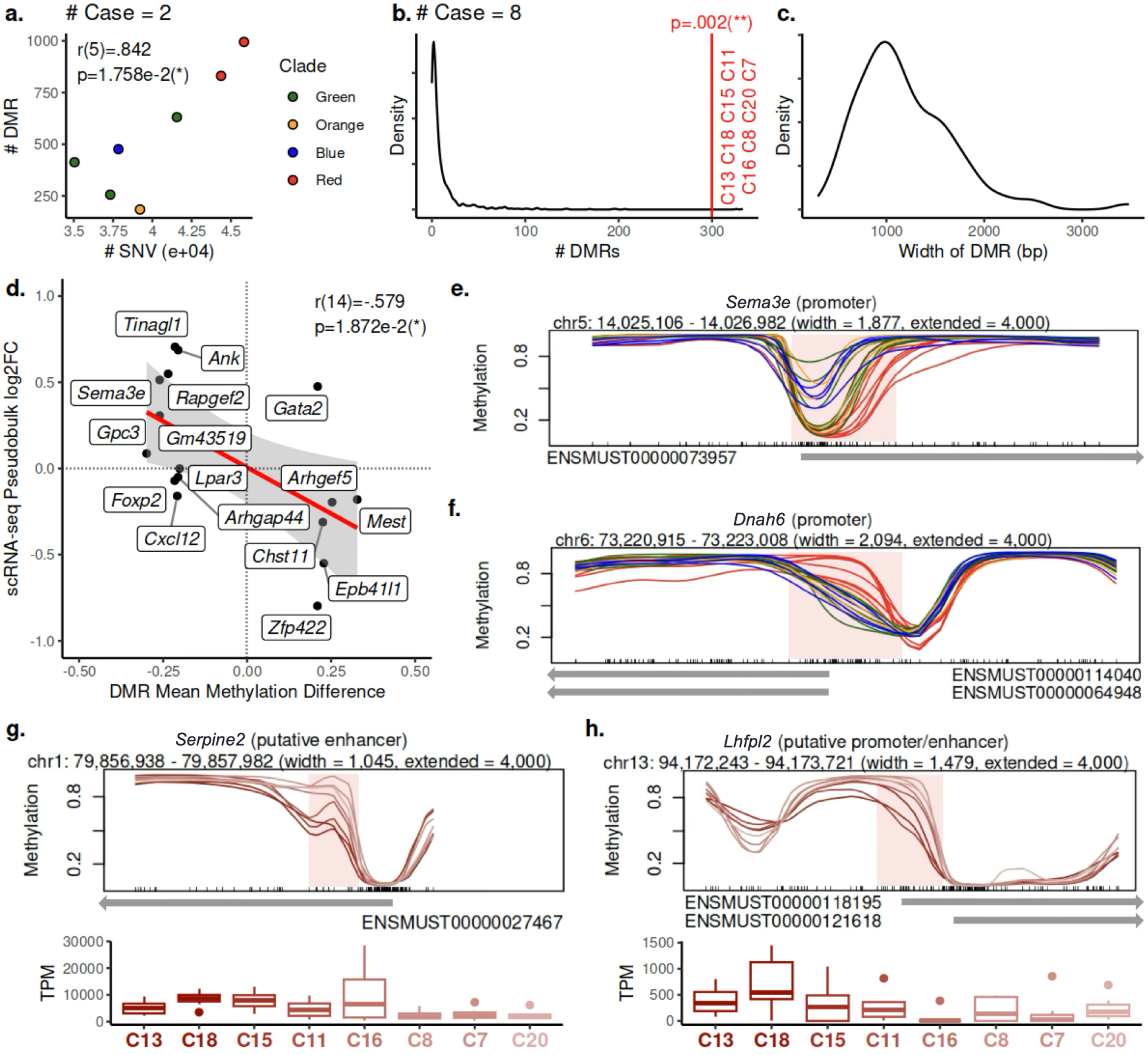
DMRs detected at subclonal resolution. **a.** The number of SNVs cumulative from trunk to the branch and the number of DMRs detected at the branch, for each branch parental to 2 leaves. **b.** The number of DMRs detected at the branch rooting the red clade and the corresponding null distribution of the number of DMRs generated during the permutation test for case groups of size 8. **c.** The distribution of the widths of the 300 DMRs detected at the branch rooting the red clade. **d.** Mean methylation difference within DMRs between sublines in the red clade and others and the corresponding log2 fold change in gene expression in genes whose (putative) promoter or enhancer the DMRs overlap as measured by scRNA-seq. **e.** DMR overlapping the promoter of *Sema3e* where sublines in the red clade are hypomethylated. **f.** DMR overlapping the promoter of Dnah6 where sublines in the red clade are hypermethylated. **g.** Genomic region overlapping a putative enhancer of *Serpine2* where it is monotonically becoming hypermethylated in the red clade while the gene expression of *Serpine2* follows a monotonicly decreasing trend. **h.** Genomic region overlapping a putative promoter of *Lhfpl2* where it is monotonically becoming hypermethylated in the red clade while the gene expression of *Lhfpl2* follows a monotonicly decreasing trend. In **e.** and **f.**, the methylation level in each subline is plotted with the color corresponding to the clade of the subline in the tumor phylogeny. In **g.** and **h.**, the methylation level and TPM values as profiled by scRNA-seq in each subline are plotted with the color hue corresponding to its phylogenetic order in the red clade.

We then focused on epigenomic analysis of the red clade since it was characterized with significantly worse survival, compared to other clades (Kaplan-Meier, *p < .*0005; Figure 2a.). For the root branch of the red clade, BSmooth [37] obtained 300 DMRs, which is significantly more than what is expected for any group of eight sublines (*p* = .002, Methods, Figure 5b.). The median length of the DMR regions was 1,081bp (Figure 5c.). Among all 300 DMRs, 17 overlap with the putative promoter regions in GENCODE transcription start sites [65, 28], and nine were annotated in the EPDnew (v.003) database of experimentally validated promoters [22, 73]. Another nine DMRs displayed enhancer-like signature and were within 2Kbp of an annotated transcription start site [25, 64, 41, 1]. (Table S5).

Notably, the EPDnew-validated promoter of *Sema3e* was found to be hypomethylated (Figure 5e.), and its human homolog has been shown to drive cancer cell invasiveness [7]. Additionally, the EPDnew-validated promoter of *Dnah6*, previously implicated in ICB response in human melanoma [101], was found to be hyper-methylated in the red clade (Figure 5f.). Coincidentally, *Dnah6* also harbors a clonal truncating mutation p.W1807* (Figure 2a.). Upon closer examination, we found evidence for haplotype-specific methylation of *Dnah6* promoter on the mutated copy in green, orange and blue clades, and the hypermethylation in the red clade was likely associated with the loss of the wild-type copy (Supplementary Figure S19).

While the red clade was distinct the rest of the phylogeny in terms of survival, it also showed the gradual changes of genotype and phenotype within the clade. The 8 red sublines followed a linear differentiation pattern, instead of a branching pattern in other clades. Further, survival monotonically improved along the evolutionary trajectory, *τ* (5)=1, *p < .*001 (Methods, Figure 2a.). To evaluate if methylation was driving gradual change in phenotype, we analyzed DMRs within the red clade along the trajectory and focused on (putative) promoter or enhancers of known genes.

We found 146 genes whose (putative) promoter or enhancer overlap with regions displaying monotonic change in methylation rate along the evolutionary trajectory, which is found to be significantly more than what is expected by chance via a permutation test on the order of the sublines within the red clade (*p* = .021, Methods). Some of these methylation changes were correlated with changes in scRNA-seq expression. In particular, a putative enhancer of *Serpine2* and the putative promoter and enhancer of *Lhfpl2* are gradually hypermethylated in the red clade, and their expression correspondingly decreases (Figure 5g.,h.). In human melanoma, SerpinE2 expression has previously been associated with high invasive potential of slow-cycling cells [83], and *LHFPL2* expression was associated with mTORC1 signaling [9], which promotes mesenchymal-epithelial transition in cancer [89].

### Systematic analysis of multiple variation types provides insights into B2905 lineages differentiation

Overall, our long-read approach offers us a unique lens to observe the complete spectrum of genomic and epigenomic events at the subclonal level and provided insights into the evolution of the B2905 melanoma (Figure 6). We detected clonal mutations of all types that affected genes known to contribute to melanoma initiation, including hotspot SNVs in *Kras* and *Gnaq*, SNVs in frequently mutated sites of *Sf3b1* and *Tfap2a*, homozygous deletion of *Cdkn2a*, and amplifications of *Met*, *Braf*, *Mitf*, *Kras*, *Cdk4*, and *Erbb3*. We observed a substantial proportion of UV damage signature on the clonal level, but not on sublonal levels; confirming UV damage as a key initiation event [103], but unlikely the driver of lineage differentiation.

**Figure 6:**
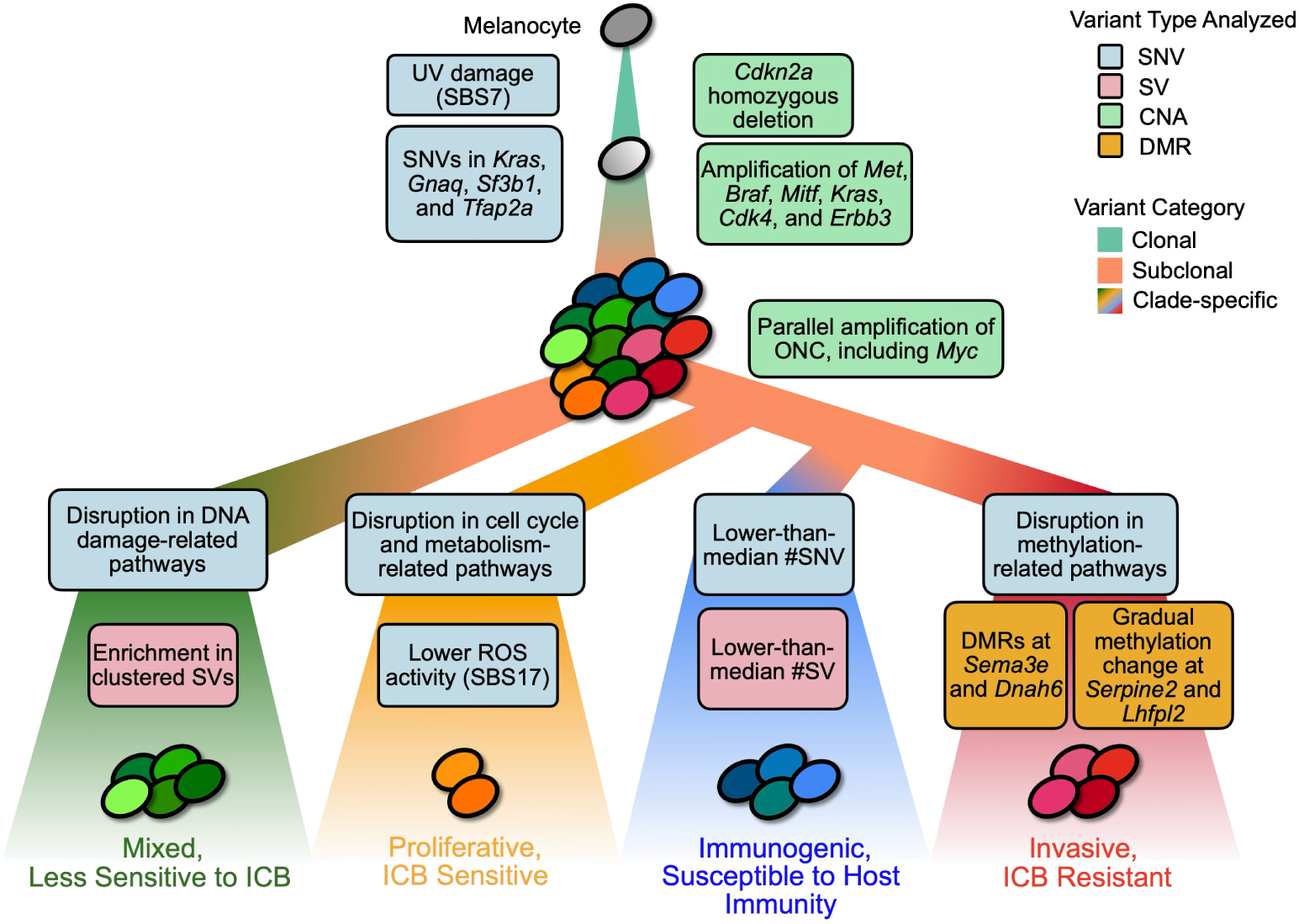
A model of B2905 melanoma evolution informed by analyses of long-read sequencing data on the 23 sublines. Key clonal, subclonal, and clade-specific observations from analyses of the SNVs, SVs, CNAs, and DMRs are labeled along the tumor evolutionary trajectories.

**Figure 7:**
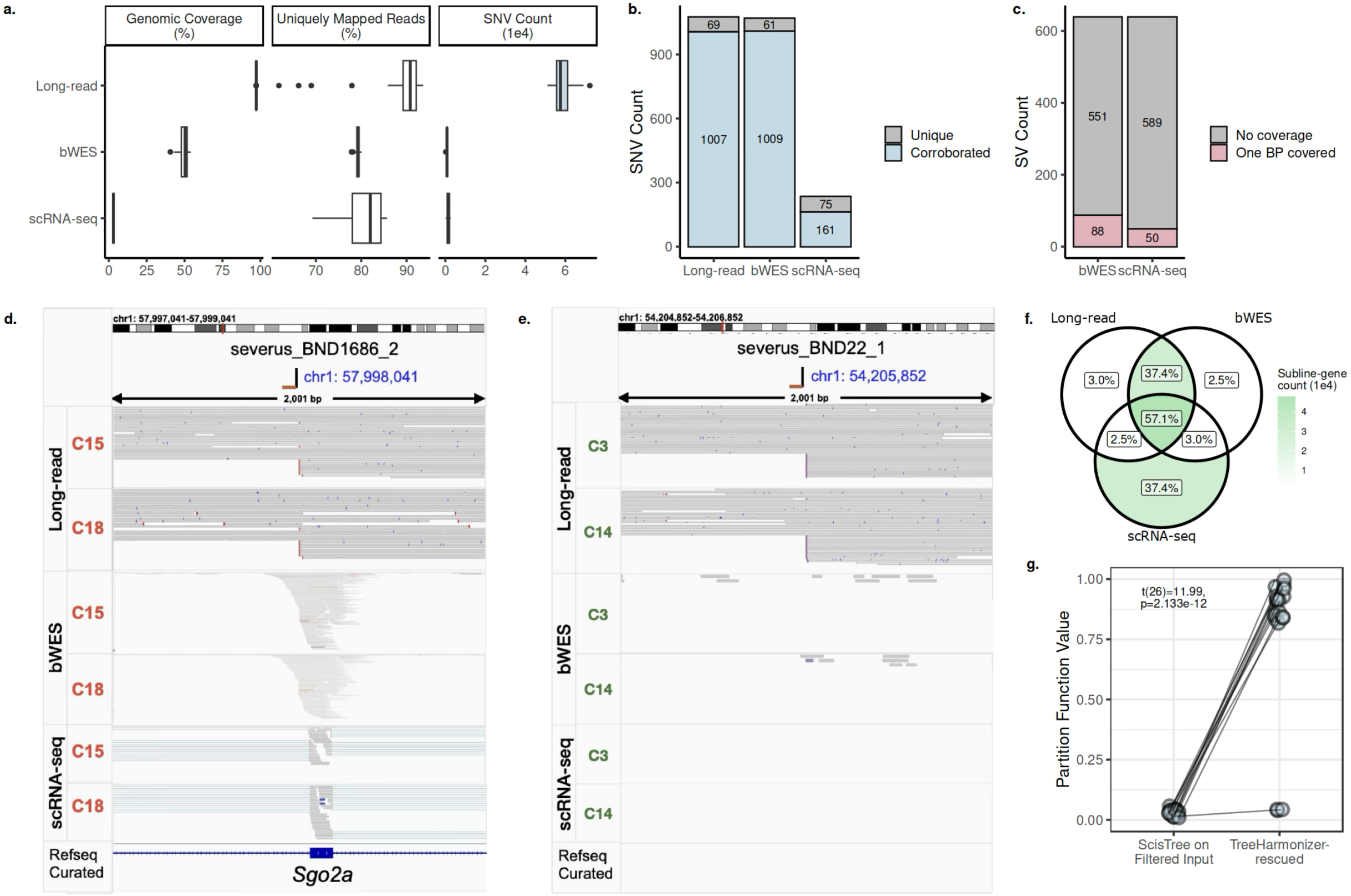
Comparison among long-read, bWES, and scRNA-seq data of the 23 sublines. **a.** Per- subline autosomal genomic coverage (%), uniquely mapped reads (%), and count of unique somatic SNVs profiled (Methods). **b.** Counts of unique, non-private somatic SNVs called within bWES target regions across the 23 sublines for each technology. The counts are stratified by whether the variants’ presence were corroborated by at least one other technology. **c.** Count of Severus-profiled SVs whose breakpoints have sufficient read coverage (200x after aggregating all reads from all sublines/cells) in bWES and scRNA-seq data. **d.** Example of genomic coverage at a near-exonic breakpoint associated with a subclonal SV across the 3 technologies. **e.** Example of genomic coverage at an intergenic breakpoint associated with a subclonal SV across the 3 technologies. **f.** Agreement on gene copy-number status (neutral, gain, loss) called from data from the 3 technologies across all sublines. For any subline, only genes with copy number calls available in all 3 technologies are included. **g.** Improvement in terms of partition function values [71] among variants impacted by filtering during tree construction, following variant placement by TreeHarmonizer.

On the subclonal level, accurate and precise SV breakpoints detection, enabled by long-read sequencing, allowed us to resolve parallel CNAs events that affected overlapping genomic regions. For example, *Myc* was independently amplified across different lineages via distinct genomic rearrangements, ranging from whole-chromosome duplications and a small, focal amplification. Overall, the positioning of CNA breakpoints relative to neighboring ONC and TSG genes suggested that these events occurred randomly, and then were independently selected.

Furthermore, our phylogenetic approach enabled the comparisons among different evolutionary trajectories of major clades that resulted in observed phenotypic heterogeneity. In the green clade, which had a significantly higher number of clustered SVs, we observed a disproportional disruption of pathways related to DNA damage signaling and response (Supplementary Figure S7c.), possibly causing an increase in genomic instability. In orange clade, the disrupted pathways related to cell cycle and metabolism and lower ROS activity cohered with its proliferative phenotype (Supplementary Figure S7c.,S8). Blue clade sublines had significantly fewer number of clade specific SNVs (one-tailed *t*-test, *p* = 3.31*e* − 02) and SVs (one-tailed *t*-test, *p* = 1.65*e* − 02); which may reflect lower selection pressure on the blue sublines, but requires further investigation.

The red clade trajectory – showing faster growth in immunocompetent hosts and higher ICB resistance – stood out in terms of increased number of characteristic DMRs, including promoters of *Sema3e* and *Dnah6* that could drive the aggressive phenotype. Interestingly, methylation-related pathways in the red clade were disproportionally impacted by non-synonymous SNVs, which may explain the increase in observed methylation changes, compared to other clades (Supplementary Figure S7f.). Moreover, we observed that a significant number of genes in the red clade, such as *Serpine2* and *Lhfpl2*, displayed a monotonic change in methylation at (putative) promoters or enhancers, often correlated with aggressiveness and scRNA-seq expression changes. This suggests that DMRs may drive a gradual phenotypic trajectory change. Overall, this showcases how joint analysis of SNVs, SVs, CNAs and DMRs provides explanation of lineage-specific phenotypes and genotypes.

### Long-read whole-genome sequencing substantially improves resolution of genomic variants

Given the complex variational landscape of the B2905 evolution presented above, we next sought to quantify the improvement in resolution enabled by long-read sequencing, as compared to the short-read bWES and scRNA-seq of B2905 subclones [40, 71].

Overall, long-read data had substantially better mappability and covered more variants (Figure 7a.), consistent with other benchmarks [58]. For our purpose of studying tumor evolution, the advantage is two-fold. First, long-read sequencing offers signals across the whole genome instead of focusing on exonic, expressed regions. This leads to (i) a substantially higher number of profiled somatic SNVs (Figure 7a.), (ii) identification of non-exonic SV breakpoints, and (iii) more contiguous estimation of copy number including non-exonic regions. Secondly, the reduction in mapping ambiguity avoids noisy signals due to mapping artifacts, amplifying real signals.

Using the available bWES and scRNA-seq data, we also validated the accuracies of the long-read profiled variants, quantifying the improvement in resolution when possible. Within bWES target regions, non-private somatic SNVs are *>* 93% consistent between long-read and bWES (Figure 7b.). Among those, *>* 15% are also corroborated by the scRNA-seq data . For each of the 639 Severus-profiled somatic SVs in long-read, we examined whether at least one of the breakends it is associated with has sufficient read coverage (200x after aggregating all reads from all sublines/cells) in bWES and scRNA-seq, as an overestimation of whether it can possibly be profiled in those technologies. Respectively, *>* 86% and *>* 92% of Severus-profiled somatic SVs have no breakends of sufficient coverage in bWES and scRNA-seq data (Figure 7c.), suggesting most SVs would have been missed in short-read data. We also profiled copy numbers from bWES and scRNA-seq data (Methods) to gauge the accuracy of long-read calls. Looking across all sublines, among genes with copy number estimates available across all 3 technologies, *>* 94% had consistent copy number status (neutral, gain, loss) between long-read and bWES (Figure 7f.). Among those, a majority also has its status corroborated in scRNA-seq calls, even though scRNA-seq calls suffer from noise and sparsity (Figure 7f., Supplementary Figure S20).

Importantly, as the detection of parallel CNA events relies on the association of contiguous copy number changes with exact SV breakpoints, using long-read whole-genome sequencing is in practice necessary for that purpose. For example, the 2 key subclonal parallel CNAs observed on chr1 (Figure 4) in our work, distinguished by their respective breakpoints, both have insufficient support in bWES and scRNA-seq data at the breakpoint loci, regardless of whether it is near-exonic (Figure 7d.) or intergenic (Figure 7e.).

In addition, we validated the TreeHarmonizer placement of SNVs using the orthogonal partition function approach [71] that estimates the likelihood of mutation placement (Methods). For the SNVs that were rescued by TreeHarmonizer, the partition function reported significantly higher scores (one-tailed paired-sample *t*- test t(26) = 11.99*, p* = 2.133*e* − 12), indicating accurate placement in the phylogeny (Figure 7g).

Aside from providing higher resolution SNV, SV, and CNA profiling, as well as being critical for identifying parallel CNAs, long-read sequencing additionally enables simultaneous profiling of methylation, which is impossible with short-read technologies without additional bisultifte-treated matched samples. Together, these demonstrate that in the specific context of studying tumor evolution, long-read whole-genome sequencing provides unrivaled advantages in revealing both genomic and epignomic insights compared to short-read alternatives.

### Curated set of subclonal variant calls enables benchmarking of long-read tools

In addition to enabling a highly detailed view of melanoma evolution, our approach provides a unique resource for development of new cancer genomics methods. Variant placement using TreeHarmonizer resulted in a highly accurate set of SNV, SV and CNA calls with established phylogenetic relationship. This enables modeling a heterogeneous tumor sample by mixing subsets of sublines sequencing data, as for each such subset a benchmarking set of variants could be easily derived. As a proof of concept, we generated 5 pseudobulk samples by merging reads from different clade combinations (Figure 8a.), and then compared several recently developed state-of-the-art tools for long-read variant calling in terms of their precision and recall on somatic variants (Figure 8b.; Methods).

**Figure 8:**
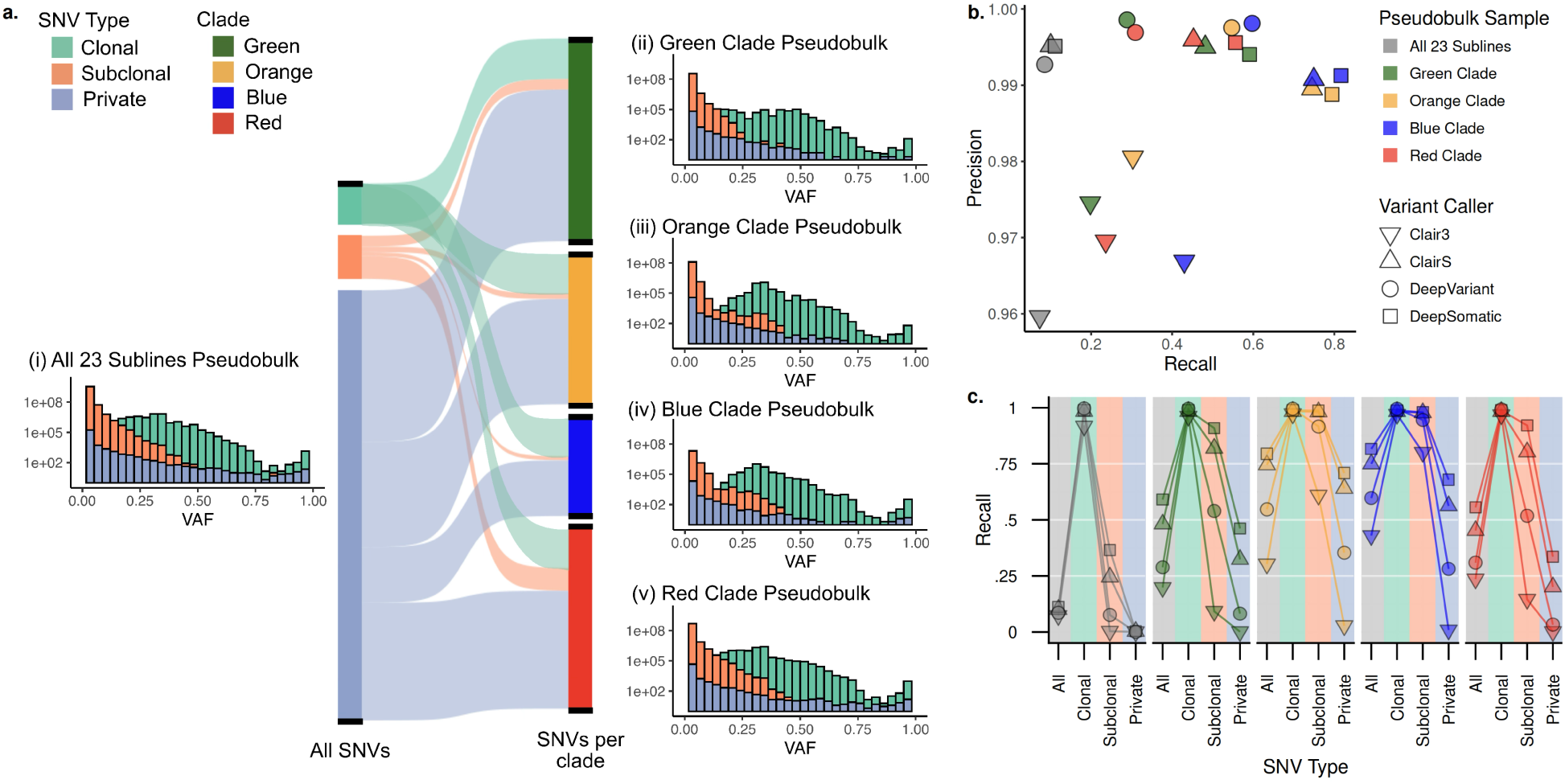
Subline-derived pseudobulk samples enables benchmarking the performances of long-read variant callers at various clonality. **a.** Five pseudobulk samples generated by merging reads from (i) all 23 sublines, and sublines in (ii) green clade, (iii) orange clade, (iv) blue clade, and (v) red clade. For each pseudobulk sample, the makeup in terms of clonal, subclonal, and private variants is shown via the Sankey diagram, and the VAF distribution of the corresponding ground truth set of variants is shown respectively. **b.** Precision and recall performance of Clair3, ClairS, DeepVariant, and DeepSomatic on the five pseudobulk samples. **c.** Recall performance of variant callers on the five pseudobulk samples, stratified by the clonality of the target variants in the ground truth set.

While precision was generally high for all callers, recall varied highly between callers and across different pseudobulk samples (Figure 8b.). Specifically, as compared to clonal SNVs, recall for subclonal and especially private SNVs was substantially lower, likely due to the decreased allelic fraction of such variants (Figure 8a.,c.). Overall, we found that recent tools that were developed for somatic analysis (ClairS and DeepSomatic) performed better than general-purpose variant callers, with DeepSomatic performing the best, maintaining the high precision as other callers while notably outperforming in recall (Figure 8b.).

This analysis underscores the difficulty of detecting rare variants from heterogeneous cancer samples – one of the key challenges in precision oncology. Even though somatic variant callers demonstrated an improvement over germline callers, even the best performers left many variants undetected in the most complex pseudobulk dataset. We publicly release the data and variant calls to encourage the development and continuous improvement of new cancer genomics methods.

## Discussion

In this study, we present an approach to comprehensively map genomic and epigenomic alterations that drive the initiation, progression and clonal differentiation of a mouse melanoma tumor model. We used Nanopore sequencing of clonal sublines – each derived from a single cell of the parental tumor – to systematically catalog somatic SNVs, structural variants (SVs), copy number alterations (CNAs), and differentially methylated regions (DMRs). We developed an algorithmic framework to place all variant types in the context of the sublines phylogeny, representing dynamics of various mutational processes along the evolutionary timescale. This approach provided an highly detailed and comprehensive view of melanoma evolution, illuminating the complex interplay between mutational processes that drive phenotypic differentiation of clonal lineages. We also demonstrated that our data can serve as a useful resource for new cancer genomics developments.

Long-read sequencing of individual sublines enabled precise resolution of parallel evolution events [67], which has been challenging to resolve using conventional approaches. Prior studies have reported examples of parallel CNAs and SVs using bulk sequencing of multiple matching samples, for example, in ovarian [69], breast [107], and osteosarcoma [50] cancers. However, these analyses were typically constrained by heterogeneity of individual samples and limited spatial or temporal resolution, making it difficult to confidently identify independent events affecting the same genomic regions. More recent efforts using single-cell DNA sequencing reported large parallel CNAs in cell line models [30] and in acute myeloid leukemia with complex karyotype [53]. Yet, due to data sparsity, these approaches often rely on pseudo-bulk inference of structural breakpoints, which prevents assignment of CNAs and SVs to individual cells. In contrast, our approach provides a highly resolved view of parallel clonal evolution thanks to the high purity of sublines, higher number of samples and the advantages of long-read sequencing.

In addition to genetic variation, Nanopore sequencing enabled the detection of lineage-specific differentially methylated regions (DMRs), offering insights into the epigenetic landscape of tumor evolution. DNA methylation has increasingly been recognized as both a marker of clonal lineage [31, 8, 60, 11] and a regulatory mechanism driving phenotypic diversity [38, 46, 10, 110]. Our work focuses on the latter, extending epigenetic resolution to the subclonal level—beyond traditional comparisons of tumor versus normal tissue or across tumor subtypes.

Recent long-read studies have explored allele-specific methylation (ASM), for example in brain tissue [51] or medulloblastoma [87]. While genome-wide ASM analysis was difficult due to the low heterozygosity of the B2905 mouse model, we illustrated that local analysis was possible in the presence of nearby somatic SNVs; furthermore, the analytical framework presented here could be readily adapted to genome-wide allele-specific analyses, for example in human.

Overall, our study highlights the advantage of long-read sequencing in resolving the the complete variational landscape of tumors, potentially benefiting precision oncology studies, as for various cancers it has been shown that variants other than SNVs, including SVs and DMRs, are important biomarkers for prognosis or therapeutic response [49, 27]. Our analytical framework, including the TreeHarmonizer tool, could be adopted to multi-site or longitudinal human tumor datasets. Finally, by publicly releasing all sequencing data and curated variant call sets, we provide a unique benchmarking resource to accelerate the development and validation of new cancer genomics tools.

## Methods

### Nanopore sequencing of B2905 sublines

Here we sequenced 23 mouse melanoma cell lines derived in a previous study [34] and 1 mouse spleen using Oxford Nanopore. gDNA was extracted using high molecular weight DNA extraction kit for cells and blood (New England Lab cat# T3050), following the manufacturer’s instruction. The extracted gDNA was further processed for nanopore sequencing following the protocol described previously [44]. Briefly, purified HMW gDNA from the 22 samples were hand-shear 20X with 1 mL Luer-Lock syringe (Becton Dickinson cat#309628) and a 0.8mm x 40 mm needle (Becton Dickinson cat#305167). Molecular DNA size was determined using TapeStation 4200 (Agilent cat#G2991BA); Most of the samples were over 55 Kb. After quantification using the Qubit dsDNA BR assay kit (Invitrogen cat#Q32853) at least 5 ug of DNA for each sample was size selected with the PacBio short read eliminator kit (PacBio cat# 102-208-300) to reduce fragments of DNA under 25 kb. After eluting the samples with 150 ul of EB buffer, DNA was quantified again by Qubit BR assay. Samples with similar concentrations (typically at 30-60 ng/ul range) were sheared simultaneously to a target size of 30 kb with the Megaruptor 3 instrument (Diagenode cat#B06010003) and the DNAFluid+ Kit for viscous DNA (Diagenode cat#E07020001) for 2 rounds at speed 45. Once the shearing was completed, samples volumes were reduced from 150 ul to 50 ul, using the speed vacuum instrument Vacufuge plus (Eppendorf cat#E-VPVCCS). The molecular DNA size was examined again using TapeStation to confirm that the samples were high molecular weight DNA. For ONT sequencing library construction, 48 ul of each sample with at least 2 ug of DNA were prepared using the Nanopore Ligation Sequencing Kit V14 (Nanopore cat#SQK-LSK114) with few modifications: (i) During DNA repair and end-prep step, 75% ethanol was used for washing instead of the recommended 70%. (ii) SFB buffer was used in the adapter ligation and clean-up step. (iii) The library elution conditions were changed to 20 min at 37*°*C. Finally, the libraries were loaded into flow cells R10.4.1 (Nanopore cat#FLO-PRO114M) for sequencing with PromethION devices. To obtain high depth coverage, each library was reloaded several times into the same flow cell, after performing a nuclease flush using the Nanopore flow cell wash kit (Nanopore cat#EXP-WSH004).

Sublines C1, C8, C11, C13, C9, C17 and the splenocyte normal were sequenced at 4khz, while the remaining sublines and the melanocyte were sequenced at 5khz, due to the frequency and requirement of ONT MinKNOW sequencing software updates. Basecalling was performed with either guppy(v6.4.2) [79] for the 4khz samples or dorado(v0.3.4) [80] for the 5khz samples. Quality of the sequencing was verified by the sequencing teams and confirmed using samtools stats [18], seqkit stats [92] and pycoQC [52]. No significant batch effects were observed between basecaller versions and sequencing frequency differences. Full sequencing statistics compiled from seqkit and pycoQC per subline are available in Table S1.

We note that we did not sequence C2 subline with Nanopore, as it has showed evidence to contamination with C3.

### Data preparation and somatic SNV calling

Long-reads were aligned using minimap2 [56] to the GRCm38 (mm10) reference. Aligned BAM files were phased and haplotagged using margin [91] and Whatshap [66] with the splenocyte normal. SNVs were called independently per subline using DeepVariant(v1.6.1) [86] with the default parameters for ONT R10.4 chemistry. Somatic SNVs were determined by subtracting both the splenocyte normal SNVs and the SNVs present in the FVB mouse line. We note that DeepSomatic [82] – a specialized long-read somatic SNV caller – became available after the majority of this analysis was completed. However, because of very few germline mutations (mm10 reference was derived from the same C57B/6J strain) and low heterogeneity within each subline, we expect DeepVariant to perform similarly to DeepSomatic on this dataset, when calling per individual subline.

SNVs for FVB were called using dipcall [57], comparing the refrence assembly for C57BL/6J to the latest long read-based FVB strain assembly [61]. FVB alignments covered most, but not all, of the C57BL/6J reference assembly, indicating either highly divergent regions or assembly errors. Since within such regions we could not distinguish somatic variants in the sublines from germline FVB variants, we removed the regions from the further analysis. Excluding those regions removed approximately 11.62% of the genome (autosome only) which in turn excluded around 14.93% of all called somatic SNVs.

### Extending Severus for multi-sample calling

SVs were called across all sublines in a single run using the multi-sample mode of Severus [48], which eliminates the need for ad-hoc re-genotyping. Unphased BAM files from each subline and the normal sample were provided as input. To account for batch effects arising from differences in N50 and read quality across samples, the --low-quality parameter was enabled. Germline SVs from the FVB strain were filtered via a panel of normals (--PON) option. FVB SV calls were generated from the FVB assembly using hapdiff [51]. Additionally, confident regions from the FVB assembly were used to define reliable genomic intervals and further reduce potential false positives.

### Copy number variant calling

We used Wakhan [2] in unphased mode to call copy number aberrations in each subline separately. As an input, Wakhan requires BAM files to extract bin level coverage and tumor phased VCFs to detect LOH. We provided Wakhan with haplotagged BAM files and the somatic variant callset VCFs per subline. We ran Wakhan with a bin size of 50kb to extract coverage from BAM file histograms. Wakhan was provided Severus generated structural variations/breakpoints VCF files so that it could detect more precise copy number change boundaries. We used an additional option, --cpd-internal-segments, to enable a change point detection algorithm that can detect small copy number changes in the rare cases when Severus misses a breakpoint.

Some genomic areas were challenging due to the poor mapping quality, reference artifacts, ends of chromosomes, and very small spans between breakpoints. However, the vast majority of such challenging regions were short. As Wakhan is optimized for calls larger than ∼50Kbp, one should take care when using calls below this threshold. We only consider calls larger than 50kb for any functional analysis in this work, unless they fall between a pair of Severus breakpoints representing one SV. We also use smaller calls for conservative region filtering during the initial tree construction, as discussed in the section below.

### Constructing tumor phylogeny using long read sequencing data from 23 sublines

To infer binary cell-linage tree of 23 sublines using ScisTree [104], we obtained a subset of highly reliable SNVs. For that, we first filtered out (i) SNVs in regions that were impacted by copy number losses in at least 1 subline, as such mutations are subject to the violations of the infinite-sites assumption and (ii) private (present in only one subline) or clonal (present in all sublines) SNVs, as these are not informative for phylogenetic analysis. We note that although some recently developed methods allow mutational losses [23, 12, 90], our filtered SNV set was large enough for reliable tree construciton using traditional methods. After that, for each mutation in each subline, we set the mutation status to “missing” if the coverage at the site is ≤ 10, deferring to the tree building algorithm to determine the presence-absence status of the mutation in the subline due to insufficient coverage. To avoid sites with consistently low coverage, as their mutation calls are unreliable across many sublines, we also filtered out sites with mean or median coverage across all sublines *<* 20. In addition, we also removed mutations with mean or median VAF across all sublines in which they were called as present below 1*/*3, a value lower than the expected VAF for most non-private heterozygous SNVs in our dataset (which is typically 0.50 for mutations from diploid regions). Lastly, we observed that some SNVs had some, but low, read support for the alternate allele in a significant fraction of sublines for which they were called as absent. Such SNVs are typically unreliable and require the tree building algorithm to make many false positive/negative corrections. We suspect that such ambiguous SNV calls are the result of various sequencing or mapping artifacts. To reduce the impact of such calls, we ignored sites for which there was read support for the alternate allele in at least half of the sublines in which the SNV was called absent. After the above filtering steps, we were left with 10,862 high-quality, phylogenetically informative somatic SNVs that were used for tree inference.

To construct the phylogenetic cell lineage tree from the filtered set of SNVs, we used ScisTree [104], a neighbor-joining based method that performs search in the space of cell-lineage trees. It is suitable for our dataset which has significantly lower number of cells compared to the number of SNVs. We performed a grid search over a range of false positive (10*^−^*^2^, 5 · 10*^−^*^2^, 10*^−^*^3^, 10*^−^*^4^, 10*^−^*^5^, 10*^−^*^6^, 10*^−^*^7^, 10*^−^*^8^) and a range of false negative (.05, .075, .1, .125, .15, .175, .2) priors. In addition, as ScisTree performs a heuristic search, its output can depend on the order of the rows (cells) in the input. Therefore, we also repeated the tree building process for 100 different permutations of the input rows. As the final output, we selected the tree with the maximum likelihood among all the 8 (false positive priors) × 7 (false negative priors) × 100 (permutations of rows) = 5600 trees inferred by ScisTree.

### Determining the phenotypes of the 4 major clades

Here we provide a brief summary of the previous work that characterized the phenotypes of the B2905 subclones [34]. After generation from parental B2905 melanoma cell line, the work performed the following characterization studies: (1) each subline was subjected to exome sequencing and scRNA analysis; (2) each subline was tested for *in vivo* growth in C57BL/6 mice; (3) selected sublines were tested for response to ICB in C57BL/6 mice.

As defined by the criteria by Hirsch et al. [40], invasive phenotype exhibits faster tumor growth when implanted in immunocompetent host, lower expression of cell cycle genes, and expression of non-canonical Wnt signaling genes; meanwhile, proliferative phenotype exhibits slower tumor growth when implanted in immunocompetent host, higher expression of cell cycle genes, and expression of canonical Wnt signaling signature. Given the relevant results from the characterization study [34], accordingly, the orange clade was determined to be proliferative, the red clade invasive, and the green clade exhibiting intermediate levels of these features was defined as having a mixed phenotype (Supplementary Table S2).

The response to host immunity (immunogenicity) of each subline was estimated by the delay of growth initiation (DGI, the time from implantation of cancer cells to starting continuous growth without regression) in immunocompetent C57BL/6 mice implanted with each subline [34]. The longer period of DGI indicates stronger response to host immunity. DGI diminished when cancer cells were implanted to immunocompromised host, as compared to that in immunocompetent host, confirming the association of DGI with tumor response to host immunity (Supplementary Figure S1).

To test the response to ICB, one or two sublines were selected from green, orange, and red clade [34]. The blue clade was not tested because of low tumor take rate (the odds that implanted cancer cells can grow tumor in the hosts). For each selected subline, its cells were implanted into ten C57BL/6, which were randomized to received anti-CTLA4 or isotype control antibody treatment. Taken together, these results of the characterization studies label each major clade with a distinct phenotype (Supplementary Table S2).

### TreeHarmonizer algorithm

Given a phylogenetic tree, TreeHarmonizer attempts to place variants on their appropriate position within the evolutionary context. Tree branches could be of following classes: truncal (occurring jointly prior to clone differentiation in this context), internal (an informative variant for clone differentiation), or terminal (a privately obtained mutation for this specific clone). Variant placement is challenging because variants of different types may overlap (e.g. SNVs inside subclonal deletion) or be miscalled in a subset of sublines.

TreeHarmonizer takes several input parameters. By default, it takes a phylogenetic tree; SNV, SV and CNA call sets; an estimated variant call FN rate; an estimated variant call FP rate; a deletion set; and a loss set. For our data, the estimated FN rate is 15%, and estimated FP rate is 0.001%, as determined by the phylogeny creation in the previous section. The SNVs call set are DeepVariant called somatic SNVs per subline. The deletion set are SVs called as deletions by Severus. The loss set are regions labeled as CN=0 or CN=1 by Wakhan.

For each distinct SNV, defined by chromosome, position, reference allele, and alternate allele, TreeHarmonizer begins by calculating its most recent common ancestor (MRCA) given the sublines in which it was called. Given the low *a priori* FP rate (0.001%) for our input, we assume that the existing variant set used to compute the MRCA is correct, and do not consider possible permutations of variant status for sublines (if the FP rate warranted it, TreeHarmonizer could consider alternate MRCAs by leaving out samples persuant to the FP rate). Given our estimated FN rate input parameter, and phylogenetic tree that is expected to consist of multiple major taxa, we define a clade-size based threshold. For clades of size 1 or 2, a variant must have the support of all sublines. For clades of size 3 and higher, the minimal subline support is the largest integer such that (*clade size* − *support*)*/clade size >*= *FN*, so that we approach as close as possible (but do not go below) the target FN rate. We allowed this to apply to clades of size 3 (even though this leads to an estimated FN rate of 33%), as the sublines within each of those clades had low branch support and high variability between the nearly-optimal phylogeny candidates.

Subline genomes may contain large deletions and structural variations that result in large losses across the genome. Such losses can lead to SNVs in those regions being deleted, which leads to an apparent contradiction with the phylogenetic tree structure under the infinite sites assumption. TreeHarmonizer models the loss of such variants, given deletion and information and assigns the “lost variant” genotype to the respective sublines. We refer to this procedure as regenotyping. In this dataset, we used SVs identified as deletions by Severus, and CNAs labeled as copy numbers 1 or 0 as regions of loss (since the majority of the chromosomes had 2 copies). Although different SV processes can lead to different loss types, all losses are treated in the same manner for regenotyping. For each SNV position, TreeHarmonizer creates a new candidate set of sublines consisting of the original calls, plus sublines where a loss may have deleted an SNV. We then evaluate the candidate set with several parameters. We consider whether the MRCA changes, and if the clade placement threshold is met with the new set of sublines. For this data, we allow for the regenotyping of all variants that pass their clade threshold for the new subline set, with two main following restrictions in order to be conservative. To account for lack of phasing information in many spans of the genome, we employ a parsimony assumption – for an SNV position that is being regenotyped, if there exists an SNV called within the loss or deletion span being considered for the same subline, we assume that the loss occurred on the other haplotype, and we do not regenotype this SNV position. Furthermore, we do not allow for dramatic changes in variant placement (a shift from a clade low on the tree to one very high on the tree), here defined as a subline reinclusion rate of *>*= 100%, as those are significantly more unlikely. If the regenotyped candidate set is rejected, the original called set is used instead. Among the placed SNVs, 1753 (0.39%) variants are regenotyped using detected deletion regions that would otherwise be considered erroneous and discarded. Full placement percentages are available in Tables S3 and S4.

A small proportion of SNVs and SVs remained unplaced after running TreeHarmonizer. The majority of unplaced SNVs fell just below the threshold limits for clonal placement, potentially indiciative of other unconsidered errors or artifacts in variant calling. SVs were placed onto the tree simply using the same clade placement thresholding formula as for SNVs. No regenotyping was performed on SVs. Since the number of unplaced SVs was relatively small compared to SNVs, we reviewed the unplaced variants and classified them into different categories. The vast majority of the 64 unplaced SVs were either in low-mappability regions characterized by increased mismatch rates (n = 21), clustered indels (n = 17) and secondary read alignments (n = 17) or mosaic repeat expansion (n = 6).

CNAs were also placed onto the tree using TreeHarmonizer. As we are not currently associating all CNAs with specific SVs, the placement of CNAs onto the phylogeny is not as evolutionarily clear. Multiple independent events may impact the same span, and thus makes it difficult to determine unique, independent, CNA spans. In order to simplify the placement problem, we limited the definition of a gain or loss to a change from a baseline of CN=2. Ranges of such gains and losses were extracted per subline from the Wakhan output.

Beginning at the leaves of the tree, we employ the following in a reverse level-order traversal — given a range *R*_1_ that corresponds to a set of sublines *S_r_*_1_, if there exists a *R*_2_ (with corresponding set *S_r_*_2_) such that *R*_1_ and *R*_2_ overlap, a new overlap range *O*_1_ is created and placed at branch MRCA(*S_r_*_1_ ∪ *S_r_*_2_), and is associated with the set of sublines *S_r_*_1_ ∪ *S_r_*_2_. As a result, every internal (and truncal) branch now contain all overlapping ranges of their children, and exactly which sublines said overlaps are associated with. Per branch, we limit the set of overlapping spans to those whose subline sets meet or exceed the minimum subline threshold for that clade size, the same as for other variant types.

Due to employing a minimum subline threshold (not requiring perfect clade completion in order to be considered valid) overlap ranges may partly overlap with each other and be associated with slightly different sets of sublines. Thus, per branch, any ranges *O*_1_ and *O*_2_ that overlap with each other are merged into a new range *O_a_*, along with their associated subline sets, *S_a_* = *S_o_*_1_ ∪ *S_o_*_2_.

We now finalize the assignment of ranges per branch with a level-order traversal. As we traverse down the tree, as ranges are assigned we exclude any span that has already been associated with a parent branch. Ranges are treated as mutable objects — if a part of said range currently being considered was already assigned to a parent, the range is split, and the unassigned remainder is placed at the current branch.

### Assessing the effectiveness of **TreeHarmonizer** in determining the placement of variants impacted by filtering during tree construction

The partition function algorithm [71] uses a tree sampling strategy to estimate the probability that, given a set of mutation calls, a mutation should be placed at the most recent common ancestor of a clade. To measure whether TreeHarmonizer effectively re-determines the placement of variants impacted by stringent filtering criteria during tree construction, we compute for those variants the respective partition function values (i) at their placement determined by ScisTree[104], and (ii) at their placement by TreeHarmonizer, with the trees sample from the space defined by the original set of variant calls prior to filtering.

We focused on the set of variants whose placement differed following the application of TreeHarmonizer. By definition, if two variants are called in an identical set of sublines in the unfiltered data, their partition function values at a given placement in the tree will be the same if the values are estimated on the same set of sampled trees. Therefore, instead of computing partition function values per variant, we compute per group of variants that (i) were called in an identical set of sublines in the unfiltered data, (ii) shared the same placement by ScisTree, and (iii) shared the same altered placement by TreeHarmonizer. We computed the partition function values with 1000 sampled trees, 0.001 false positive rate and 0.15 false negative rate, and the results on the 27 groups are reported in (Figure 7g.). One-tailed paired-sample *t* -test was applied to the partition function values of the 27 groups.

### Mutational signature analysis along the tumor phylogeny

The cumulative placed SNV profiles of the 23 sublines were used as input samples to SigProfileAssignment (v0.1.9) [20], a mutational signature fitting tool chosen for its superior overall performance and specifically for the number of mutations per sample at hand [70]. The 23 SNV profiles were translated into mutational spectra over 96 categories for joint refitting, and COSMICv3.4 [93] mm10 signatures were used as the reference catalog, as it has been shown that excluding sets of signatures from refitting often leads to poorer performance [70].

SigProfilerAssignment outputted signature activity estimates per subline, as well as signature probability per individual mutation per subline. Each mutation placed at a particular branch was assigned the most probable signature given the aggregated signature probabilities for that mutation in sublines in the subtree below the branch. The activity of a mutational signature at a branch was then estimated by the proportion of mutations placed at the branch assigned to the signature. For concision of presentation, activities for signatures SBS7a, SBS7b, SBS7c, and SBS7c were aggregated and referred to as signature “SBS7”, and those for signatures SBS17a and SBS17b were aggregated and referred to as signature “SBS17”.

For each active signature, to assess whether signature activities have significant differences according to their evolutionary timing or between clades, each branch in the tumor phylogeny is labelled by (i) truncal, internal, terminal, and (ii) the color of the clade it is in (the trunk and two branches closest to the trunk have NA as clade assignment), and analysis of variance (ANOVA) with post-hoc Tukey HSD (Honestly Significant Difference) test was performed to see if there were significant differences between pairs along either independent variable. Since there is only 1 truncal branch, and the 3 branches closest to the root are only grouped together due to lack of clade labels, numerically significant results (adjusted *p*-value *<* .05) involving those as a group in the pair are not reported. In other words, only significant results between internal and terminal branches, or between pairs of clades are reported (Supplementary Figure S8).

### Computing dN/dS values along the tumor phylogeny

Placed SNVs were annotated using Ensembl Variant Effect Predictor (v102) [68]. For any given branch, we computed the dN/dS value as the ratio of cumulative non-synonymous SNVs placed along the path from the trunk and including the current branch to that of cumulative synonymous SNVs, normalized by the ratio of synonymous sites to non-synonymous sites [40]. dN/dS values at internal branches were found to be significantly less than those at terminal branches via a one-tailed *t*-test (Supplementary Figure S7a., *p < α* = .05).

### Pathway enrichment analysis

Given a set of genes of interest - either that they are impacted by a certain type of variant at a particular time during the tumor evolution (e.g. placed at the trunk of the phylogeny) or in a particular clade - we performed Gene Ontology over-representation analysis to examine their potential functional impact [4, 15]. Genes without a canonical name (“*Gm-*” and “*-Rik* ” genes) were excluded from analysis. Using clusterProfiler (v3.21) [108], we tested whether the candidate gene set significantly over-represents a Gene Ontology Biological Process gene set between sizes of 3 and 500. The resulting *p*-values were adjusted using the Benjamini-Hochberg procedure, and terms with *q*-value *< .*05 were determined to be statistically significant.

### Determining genes impacted by SVs

We used Padfoot (https://github.com/KolmogorovLab/Padfoot, commit ID:2d5237b) to annotate SV calls generated by Severus using the --specie mouse option and GRCm38.p4 GFF3 files with basic gene an- notations. The *possible fusion* and *exon altering* events were manually validated using IGV and scRNA-seq alignments.

### Determining the set of cancer-relevant mouse genes-of-interest according to the COSMIC Gene Census

To obtain a set of mouse genes that likely have functional impact in cancer, we selected those with one-to-one homology as genes curated in the COSMIC Cancer Gene Census (CGC, v101) [93]. Genes whose COSMIC homologs are uniquely labeled as ONC in CGC are labeled as ONC, and those whose COSMIC homologs are uniquely labeled as TSG are labeled as TSG. Those whose COSMIC homologs have no unique ONC/TSG labelings in CGC are labeled as ND (not designated). Among those without unique COSMIC ONC/TSG labelings, 4 genes (*Ezh2* : ONC [113]; *Rad21* : TSG [39]; *Tert* : ONC [35]; *Trp53* : TSG [94]) were given unique labels according to literature on melanoma or other cancer types A gene is said to be impacted by CNA in a subline if it is entirely contained in a copy number gain or loss segment.

To determine if the gain or loss of a gene may be associated with its designation as ONC or TSG, we performed two *χ*^2^ tests on the set of COSMIC ONC/TSG genes impacted by CNA in any subline. One *χ*^2^ test evaluated enrichment of ONC that were gained in some subline but not lost in any sublines. The other performed the corresponding test of TSG lost in some subline but not gained in any subline. The result was considered significant if *p < α* = .05.

To see whether selection may be at play, we further examined the distribution of ONC/TSG around CNA boundaries. To that end, we specifically focused on COSMIC homolog pairs that are adjacent to each other along the genomic coordinate, where one is labelled ONC and the other TSG and have different absolute copy numbers in any subline, as they correspond to the nearest ONC/TSG COSMIC homologs upstream/downstream to a copy number change boundary. We counted in total 14 instances where ONC homolog is on the side of the boundary with higher copy number state, and TSG homolog is on the other. To assess the statistical significance of the observation, a null distribution was generated by counting instances of such observation in 5,000 permutations of the ONC/TSG/ND labels of the COSMIC homologs. The *p*-value is the fraction of times the permutated labelings yield a greater or equal number of such observations than the actual labeling. We reject the null hypothesis if *p < α*=.05.

### Timing analysis of chr4 copy-neutral LOH

The copy number calls from Wakhan suggest whole-chromosome copy-neutral LOH on chr4. Compared to other autosomes, there is a large proportion of clonal SNVs on chr4 that are homozygous, or in other words, have a median VAF of 1 across sublines evenly distributed across the chromosome (Supplementary Figure S17a.,b.). The number of such chr4 clonal SNVs provide evidence for the approximate timing of the underlying chromosomal events. Copy-neutral LOH is likely the result of one of the following two scenarios: (i) LOH followed by whole-chromosome duplication of the remaining haplotype, and (ii) mitotic nondisjunction followed by LOH in the trisomic cell; nevertheless, the scenarios are indistinguishable given current data because neither a subline with one nor a subline with three copies of chr4 was observed.

In the first scenario, we assume the copy-neutral LOH is a result of LOH followed by duplication (Supplementary Figure S17c.). Without additional information, we cannot distinguish SNVs obtained before the LOH event and those obtained after LOH and before duplication, as in either case they would appear homozygous; that said, the number of observed homozygous SNVs upperbounds the number of SNVs obtained prior to LOH. Assuming the rate of SNV acquisition per genomic length is constant, the total number of SNVs acquired prior to whole-chromosome duplication is at most two times the number of observed homozygous SNVs. Accounting for the unobserved SNVs in the lost copy in the total number of clonal SNVs, we also add the number of observed homozygous SNVs to the total number of observed chr4 clonal SNVs. The fraction of SNVs acquired prior to whole-chromosome duplication out of total clonal SNVs gives an overestimation of the timing of the duplication, which turns out to be .53, or around half way through the trunk of the tumor phylogeny.

In the second scenario, we assume the copy-neutral LOH is a result of LOH in the trisomic cell formed following mitotic nondisjunction (Supplementary Figure S17c.). In this case, without additional information, we cannot resolve the timing of the LOH event from VAF distributions of observed clonal SNVs; however, the timing of mitotic nondijunction can be similarly estimated. The number of observed homozygous SNVs corresponds to the number of SNVs on the chr4 copy with Cdkn2a deletion in a diploid cell prior to mitotic nondisjunction. Again, assuming the rate of SNV acquisition per genomic length is constant, the total number of SNVs acquired prior to mitotic nondisjunction is roughly twice the number of observed homozygous SNVs. In the trisomic cell, the three copies of chr4 acquired SNVs independently. The number of observed number of chr4 clonal heterozygous SNVs, or in other words, those with median VAF *<* 1, corresponds to that acquired after mitotic nondisjunction, normalized such that the amount of genomic content is equivalent to two copies of chr4. The fraction of SNVs acquired prior to mitotic nondisjunction out of total SNVs acquired gives an overestimation of the timing of mitotic nondisjunction, which is again, .53.

The above analyses lead to the conclusion that, assuming the rate of SNV acquisition per genomic length is constant, the tumor cell first acquired two of the chr4 copy carrying *Cdkn2a* deletion half way through the trunk of the tumor phylogeny, at the latest.

### Subclone-specific differential methylation analysis

Given modified base calls, modified base counts for each sample are aggregated using modbam2bed [63] (--aggregate --mod base=5mC --cpg --mask). The number of methylated reads (N_mod_) and total read count (N_mod_ + N_canon_) at each CpG site with methylation calls for the 23 subline samples are compiled for smoothing using BSmooth [37].

At a given branch in the phylogeny, the 23 sublines are bipartitioned into a case group, consisting of sublines in the subtree below the branch, and a control group, consisting of the others. With the case-control group assignment of sublines, we obtain a list of regions that are differentially methylated using BSmooth [37]. To reduce noise due to low coverage, only CpG sites covered by at least 3 reads in all sublines are considered. Signal-to-noise *t*-statistics were computed per CpG site using the smoothed methylation values, and a DMR is defined to be one that spans at least 10 consecutive CpG sites, with corrected *t*-statistics *< .*025 or *> .*975, and with absolute mean differences between groups at least .2. As replicates are required for DMR testing using BSmooth, DMRs were computed at branches where the bipartition of the sublines they induce both have a size of at least 2.

### Permutation test on the number of DMRs obtained

Given a case group of a certain size, we used a permutation test to assess whether significantly more DMRs were identified in the case group than in the control group. A null distribution was generated by 1,000 (or, when the size of the case group is 2, all possible 253) random permutation of the case and control group labels over the sublines. We recorded the number of DMRs identified in each permutation using the permuted labels to implement the criteria described in the section above. The value of *p* is the fraction of permutations that yield a greater or equal number of DMRs than the actual case-control groups of interest. We reject the null hypothesis if *p < α* = .05.

### Correlation between the number of DMRs and cumulative SNVs at a branch in the phylogeny

As the numbers of DMRs are not immediately comparable between tests involving case groups of different sizes, we focused on tests involving case groups of size 2 performed at branches parental to exactly 2 leaves, since there is a sufficient number (7) of such branches to assess correlation. The Pearson correlation was assessed between the cumulative numbers of SNVs from the trunk to the branch, and the number of DMRs detected between the 2 sublines below the branch and the rest of the 21 sublinesWe reject the null hypothesis if *p < α*=.05.

### Differential gene expression analysis of subline scRNA-seq data

scRNA-seq data was obtained as previously described [40], and gene-level expected count values for the subline scRNA-seq data were obtained using RSEM [54]. As multiple cells were sequenced from each subline, the analysis treated them as biological replicates of the same subline. Genes with total count less than 2^10^ were removed, and the decimal expected count values were rounded to the closest digit. DESeq2 [62] was used to assess differential gene expression, and the shrinkage estimator uses a normal prior (coef=2). The log2 fold change values for the genes of interest were obtained. Note that not all genes of interest have their expression values available in the scRNA-seq data set, and hence they are left out of the test for correlation.

### Discovering key regions where changes in methylation is monotonic along the evolutionary trajectory

To obtain candidate DMRs to test for monotonicity, we collected regions that exhibited differential methylation among the sublines within the red clade that potentially exhibit such monotonic trend. As replicates are required for DMR testing with BSmooth, 4 DMR tests were performed with increasingly small subtrees as the case group (and the rest of the sublines in the red clade as the control group): first C11, C16, C8, C7, and C20; then C16, C8, C7, and C20; followed by C8, C7, and C20; and lastly C7 and C20. These groupings were chosen as they reflect cuts along the linear evolutionary trajectory in the red clade. The union of the 4 sets of DMRs were then taken, merging overlapping intervals. As this analysis is motivated by phenotypic observations, we focused our analysis on the regions overlapping (putative) promoters and enhancers of known genes, which we refer to as candidate DMRs.

For each of the candidate DMRs, the Mann-Kendall test was performed on four orderings of the sublines – the order in which the sublines were placed (C13, C18, C15, C11, C16, C8, C7, C20), as well as the three variations that may be formed by swapping immediate siblings (C18 and C15, C7, and C20). Kendall’s *τ* value with the greatest absolute value was recorded. We set a minimum threshold of .7 for the absolute value of Kendall’s *τ*, as under assumptions it roughly corresponds to a Pearson correlation coefficient of .9 [33], which is commonly accepted to signify a strong magnitude of effect. Genes whose (putative) promoters or enhancers overlapped regions with an absolute value of Kendall’s *τ* over the threshold .7 were kept. How the statistical significance of the discovery was assessed is discussed in the following section.

### Permutation test on the number of genes associated with monotonic methylation changes along the evolutionary trajectory

We assessed the statistical significance of the number of genes we discover to be associated with monotonic methylation changes along the evolutionary trajectory in the red clade via a permutation test. Fixing the structure of the red subtree, we permuted the position of the 8 sublines in the structure. While there are 8! total permutations of 8 sublines, given the two sets of immediate siblings that are interchangeable in terms of order, and that there is symmetry in terms of the magnitude of correlation for the exact reverse of a particular order, for every permutation, there are 7 others that are equivalent for the purpose of our analysis. We obtain 1,000 permutations from the 7! equivalence classes, sampling no more than one permutation from each equivalence class.

For each permuted positioning of 8 sublines in the structure of the red subtree, we obtained the number of genes whose (putative) promoters or enhancers overlapped regions that show monotonicity in terms of methylation following the given order using the algorithm described in the previous section. The *p*-value is the fraction of times the permuted orderings yield a greater or equal number of such genes than the actual ordering of the red clade suggested by Figure 2a. We reject the null hypothesis if *p < α* = .05.

### Assessing sequencing statistics on the long-read, bWES, and scRNA-seq data of the 23 sublines

The bWES and scRNA-seq data (several cells per subline) for 23 sublines were obtained from prior studies [71, 40]. Coverage data per technology was generated with samtools coverage [18], and a weighed average was computed over the genome from per contig results. Read quality and count statistics such as N50, Q20(%), Q30(%) statistics were generated using seqkit stats [92] and samtools view -c [18]. Uniquely mapped reads were generated using sambamba [98] with different parameters per aligner formatting. For bWES data (aligned with bwa mem [55]), unaligned, secondary, and those reads with XA or SA tags and a MAPQ of 0 were removed. For long-read data, (aligned with minimap2 [56]), unaligned, secondary and MAPQ of 0 reads were removed. For scRNA-seq data, (aligned with STAR [21]), a MAPQ of 255 defines uniquely mapped reads. For per-subline analyses of scRNA-seq data, individual sample bams were merged per subline that they correspond to. Statistics for per-subline scRNA-seq data were generated using the same procedures as per scRNA-seq sample.

### Calling somatic SNVs from bWES and Smart-Seq2 full-length scRNA-seq data on the 23 sublines

Small variants were called from bWES data per subline using DeepVariant(v1.6.1) [86] with the default parameters provided for the WES model. Called variants were filtered down to SNVs and the target exome capture region. Somatic SNVs were determined by subtracting the long-read ONT splenocyte normal SNVs and the SNVs present in the FVB mouse line, the same methodology as for the long-read sequenced sub- lines. A bWES splenocyte sample was unavailable, and for equivalent somatic variant determination and comparison, the long-read ONT sample was used.

Small variants were called from scRNA-seq data per sample using DeepVariant(v1.5.0) [86] with a custom RNA-seq model developed by the DeepVariant team [16]. It was necessary to use v1.5.0 due to DeepVariant transitioning to a different model architecture as of v1.6.0 than what the RNA-seq model was built on. Calling differences are not expected within the autosome between caller versions. Somatic SNVs were determined following the same methodology as for bWES. SNVs were filtered to bWES target regions and limited to those with a coverage of 8x or higher, and to those with a genotype quality of 18 and higher, as recommended per model benchmarking by the developers [16].

### Calling copy number from bWES and Smart-Seq2 full-length scRNA-seq data on the 23 sublines

CNVkit [97] was used to obtain the CNA calls from bWES data. CNVkit was run with external bWES data from a normal spleen sample [103] as reference. The per-bin copy numbers were shown in Supplementary Figure S20, and the per-segment copy number calls were used to determine the copy number status (i.e. loss, neutral, gain) of available genes in a subline.

Full-length scRNA-seq data were obtained using the Smart-seq2 protocol [85] for several single cells per subline [40], and inferCNV [99] was used to obtain the CNA calls from the data. As Smart-seq2 data for normal melanocytes were not generated for the same study, inferCNV was run with external Smart-seq2 data from mature melanocytes [43] as reference (cutoff=1,sd amplifier=2,noise logistic=TRUE; median filtering applied). The inferCNV i6 Hidden Markov Model was used to determine the copy number state, a posterior probability of .5 was used for filtering out likely normal observations, and a per-gene copy number status for each subline was derived.

### Benchmarking using pseudo-bulk simulations

We simulated pseudo-bulk tumor sequencing data by mixing together subset of sublines, mimicking tumor samples with various degree of heterogeneity. We created five pseudo-bulk samples, one that merges reads from all sublines into one sample, and four that merge reads from the sublines for each major color clade. For each pseudo-bulk sample, a corresponding set of variants placed by TreeHarmonizer was used as a benchmarking variant set. Variants that were unplaced by TreeHarmonizer were excluded from each call set, in order to not penalize for variants not determined to necessarily be false positives.

We evaluated precision and recall of several state-of-the-art variant callers: DeepSomatic 1.7.0 [82], ClairS 0.3.1 [112], DeepVariant 1.6.1 [86] and Clair3 1.0.10 [111]. Each caller was run with its default settings for ONT R10.4 chemistry. Somatic pseudobulk variants were determined via the same methodology as for per-subline SNV calls, see section above. ClairS showed a slight improvement when using our phasing information, haplotagged using Margin [91] and Whatshap [66] per pseudo-bulk sample (the same tools and parameters as per-subline SNV phasing) – rather than using its internal intermediate phasing step. Only this run is shown in Figure 8b.,c.. As expected, as the number of clones in a pseudo-bulk dataset increases, the calling power substantially decreases, as seen by the reduced recall.

Overall, somewhat low recall rates on heterogeneous pseudo-bulk simulations suggest that there is still room to improve the variant calling methods for cancer data. This is not surprising, as specialized methods, such as DeepSomatic and ClairS, were released only recently, in parallel with our study. Public release of our dataset and variant calls will encourage further developments. We also note that while DeepSomatic performed better than DeepVariant on pseudo-bulk, we found that DeepVariant was a good fit for variant calling on individual sublines that represent a single clone due to low intrasubline heterogeneity.

## Code availability

**TreeHarmonizer** is available at https://github.com/KolmogorovLab/TreeHarmonizer.

## Data availability

Unaligned long-read sequencing data (.bam files) are available at NCBI SRA: https://www.ncbi.nlm.nih.gov/bioproject/1307171

Variant calls and other processed data are available at: https://doi.org/10.5281/zenodo.16883901[105]

## Acknowledgments

This research was supported in part by the Intramural Research Program of the National Institutes of Health (NIH). The contributions of the NIH authors were made as part of their official duties as NIH federal employees, are in compliance with agency policy requirements, and are considered Works of the United States Government. However, the findings and conclusions presented in this paper are those of the authors and do not necessarily reflect the views of the NIH or the U.S. Department of Health and Human Services. This work utilized the computational resources of the NIH HPC Biowulf cluster (https://hpc.nih.gov).

## Conflict of interest

The authors declare no competing interests.

## Notes

### Competing Interest Statement

The authors have declared no competing interest.

