## Supplementary Information for "Long-read sequencing of single cell-derived melanoma subclones reveals divergent and parallel genomic and epigenomic evolutionary trajectories"

December 2024

Yuelin Liu*, Anton Goretsky*, Ayse Keskus, Salem Malikic, Tanveer Ahmad, E. Michael Gertz, Farid Rashidi Mehrabadi, Michael Kelly, Maria Hernandez, Charlie Seibert, Juan Manuel Caravaca, Kayla Kline, Yongmei Zhao, Ying Wu, Biraj Shrestha, Bao Tran, Arindam Ghosh, Xiwen Cui, Antonella Sassano, Laksh Malik, Breeana Baker, Cornelis Blauwendraat, Kimberley J. Billingsley, Eva Perez-Guijarro, Glenn Merlino, Erin K. Molloy, S. Cenk Sahinalp#, Chi-Ping Day#, Mikhail Kolmogorov#

#### Contents

#### List of Figures

S1 **Delay of growth initiation (DGI) of sublines in C57BL/6 and nude mice.** DGI is defined as the number of days between cell implantation in a mouse and the tumor started continuous growing without regression. **a.** One million cells of each subline were implanted into each of the five C57BL/6 mice (Day 1). The tumor size was measured twice a week. The mean DGI of the sublines in each clade, as well as the median across sublines means within a clade, was shown. No tumor was grown from any mice implanted with cells of C19, C6, or C13. Note that two mice implanted with C5 cells did not grow tumor, and they were not included in the calculation of mean DGI. These results suggested that, on average, sublines in blue clade induced the stronger host immunity than those in other clades. **b.** To investigate the association of the DGI with host immunity, the subline from each clade with the highest DGI following C57BL/6 implantation was tested in immunocompromised host. For each of C19 (green clade), C5 (orange clade), C6 (blue clade), and C13 (red clade), one million cells were implanted into each of the five immunocompromised nude mice (Day 1), and the DGI values measured from each mouse, as well as the median across mouse DGI values for a subline are shown. As expected, we observed the tumor grew in all mice with diminishing DGI as compared to the implantation in C57BL/6 mice. These results indicated that DGI is associated with subline response to host immunity. . . . . . . . . . . . . . . . . . . . . . . . . 6
S2 **SNVs and SVs placed by `TreeHarmonizer`.** **a.** Cumulative number of placed SNVs and SVs per subline, colored by clade. Medians are marked by dashed lines. **b.** The number of SNVs and the number of SVs placed at each branch of the phylogeny strongly correlate with each other. **c.** Variant allele frequency distributions of placed SNVs. **d.** Variant allele frequency distributions of placed SVs. . . . . . . . . . . . . . . . . . . . . . . . . . . . . . . . 6
S3 **SNV density and SV events present in individual sublines in the green clade.** . . . 9
S4 **SNV density and SV events present in individual sublines in the orange clade.** . . 10
S5 **SNV density and SV events present in individual sublines in the blue clade.** . . . 11

S6 **SNV density and SV events present in individual sublines in the red clade.** . . . . 12
S7 **SNVs along the tumor phylogeny. a.** dN/dS value distributions among truncal, internal, and terminal branches. dN/dS values at internal branches are significantly lower than that at terminal branches by one-tailed *t*-test. **b.** Select enrichment of genes harboring non-synonymous clonal mutations among GO Biological Process term. **c.-f.** Select enrichment of genes uniquely harboring non-synonymous mutations in the green (c.), orange (d.), blue (e.), and red (f.) clade among GO Biological Process terms. . . . . . . . . . . . . . . . . . . . . . . . . 13
S8 **Mutational signature activities along the tumor phylogeny.** In each panel, a point represents a branch in the phylogeny. A branch can either be categorized into truncal, internal or terminal, or by the color of the clade it is in. ANOVA with post-hoc Tukey HSD (Honestly Significant Difference) Test was performed, and the adjusted *p*-values of statistically significant pairs are reported. . . . . . . . . . . . . . . . . . . . . . . . . . . . . . . . . . . . . . . . . 14
S9 **SV-disrupted genes and gene fusion annotated by `Padfoot`. a.** A clonal deletion affects exon 4-9 of *Nav3*. Long-read genomic reads of an representative subline from each major clade are shown. **b.** A subclonal deletion in C3 and C14 in *Met* resulted in the loss of exon 4-11. Long-read genomic reads from sublines C3 and C14 are shown. **c.** A subclonal deletion in C3 and C14 resulted in the fusion of *Map4* and *Smarcc1*. Long-read genomic reads from sublines C3 and C14, as well as scRNA-seq reads from a C14 cell corroborating the fusion event are shown. . . . . . . . . . . . . . . . . . . . . . . . . . . . . . . . . . . . . . . . . . 15
S10 **Mouse homologs of COSMIC genes impacted by (1) non-synonymous SNV, (2) SV breakpoint, (3) CNA, or (4) (hypo/hyper-) DMRs in their (putative) enhancers or promoters in 23 sublines (1/7).** Genes with "NA" copy number in a subline span detected CNA boundaries and therefore cannot be assigned a unique copy number. . . . . . . 16
S11 **Mouse homologs of COSMIC genes impacted by (1) non-synonymous SNV, (2) SV breakpoint, (3) CNA, or (4) (hypo/hyper-) DMRs in their (putative) enhancers or promoters in 23 sublines (2/7).** Genes with "NA" copy number in a subline span detected CNA boundaries and therefore cannot be assigned a unique copy number. . . . . . . 17
S12 **Mouse homologs of COSMIC genes impacted by (1) non-synonymous SNV, (2) SV breakpoint, (3) CNA, or (4) (hypo/hyper-) DMRs in their (putative) enhancers or promoters in 23 sublines (3/7).** Genes with "NA" copy number in a subline span detected CNA boundaries and therefore cannot be assigned a unique copy number. . . . . . . 18
S13 **Mouse homologs of COSMIC genes impacted by (1) non-synonymous SNV, (2) SV breakpoint, (3) CNA, or (4) (hypo/hyper-) DMRs in their (putative) enhancers or promoters in 23 sublines (4/7).** Genes with "NA" copy number in a subline span detected CNA boundaries and therefore cannot be assigned a unique copy number. . . . . . . 19
S14 **Mouse homologs of COSMIC genes impacted by (1) non-synonymous SNV, (2) SV breakpoint, (3) CNA, or (4) (hypo/hyper-) DMRs in their (putative) enhancers or promoters in 23 sublines (5/7).** Genes with "NA" copy number in a subline span detected CNA boundaries and therefore cannot be assigned a unique copy number. . . . . . . 20
S15 **Mouse homologs of COSMIC genes impacted by (1) non-synonymous SNV, (2) SV breakpoint, (3) CNA, or (4) (hypo/hyper-) DMRs in their (putative) enhancers or promoters in 23 sublines (6/7).** Genes with "NA" copy number in a subline span detected CNA boundaries and therefore cannot be assigned a unique copy number. . . . . . . 21
S16 **Mouse homologs of COSMIC genes impacted by (1) non-synonymous SNV, (2) SV breakpoint, (3) CNA, or (4) (hypo/hyper-) DMRs in their (putative) enhancers or promoters in 23 sublines (7/7).** Genes with "NA" copy number in a subline span detected CNA boundaries and therefore cannot be assigned a unique copy number. . . . . . . 22
S17 **SNV evidence for the timing clonal CNA events. a.** Median (across sublines) VAF distribution for clonal SNVs on chr4 and other autosomes. **b.** Genomic distributions of homozygous (median VAF = 1) and heterozygous (median VAF < 1) chr4 clonal SNVs. **c.** Two indistinguishable scenarios that may have resulted in the copy-neutral LOH. . . . . . . . 23

S18 **Permutation tests on the number of DMRs obtained for all subtree sizes of the phylogeny in Figure 2a.** In each panel, "# Case" indicates the number of leaves in the subtree below the branch at which the test was performed. See Figure 5b. for "# Case = 8". Note that due to symmetry, no additional permutation test is done for "# Case = 16" to account for the subtree consisting of the orange, blue, and red in Figure 2a., as the empirical null distribution is identical to that of "# Case = 7" and the number of DMRs obtained is identical to that of the subtree of size 7 colored in green in Figure 2a. . . . . . . . . . . . . . 23

S19 **Associating truncating *Dnah6* mutation with (haplotype-specific) methylation of *Dnah6* promoter in major clades.** One representative subline from each major clade is shown. Clonal truncating mutation *Dnah6* p.W1807* is marked in the left panel displaying unphased reads, and a heterozygous variant chr6: 73,220,494 [C>T] is marked on the right panel with reads phased according to the variant. In C11, the high VAF of *Dnah6* p.W1807*, together with the homozygosity of chr6: 73,220,494 [C>T], not only indicates LOH in C11, but also provides evidence that the two variants exist on the same haplotype. As the phased reads on the right panel are also colored by DNA methylation, we observe that methylation at *Dnah6* promoter is specific to the haplotype harboring chr6: 73,220,494 [C>T], and hence *Dnah6* p.W1807*. We observe haplotype-specific methylation of the *Dnah6* promoter in the green, orange, and blue clade, and hypermethylation in the red clade. Note that while here we observe a few reads from the wild-type copy of *Dnah6* in C11, we see a complete loss of the wild-type copy of *Dnah6* in all other sublines in the red clade, and along with that a complete hypermethylation of the *Dnah6* promoter. . . . . . . . . . . . . . . . . . . . . . . . . . . . . 25

S20 **CNA profiles of 23 sublines obtained using long-read, bWES, and scRNA-seq data. a.** Long-read-derived CNA profiles using `Wakhan` [2] (Figure 2e.; Methods) **b.** CNA profiles derived by `CNVkit` [14] from bWES data [11](Methods). **c.** CNA profiles derived by `inferCNV` [15] from full-length scRNA-seq data [7] obtained using the Smart-seq2 protocol [13] (Methods). . . . . . . . . . . . . . . . . . . . . . . . . . . . . . . . . . . . . . . . 26

### List of Tables

S1 **Sequencing statistics for the 23 sublines, spleen, and parental B2905 line.** Q20(%) (and Q30(%)) represents the percentage of bases that have a PHRED quality score of 20 (and 30) or over. Avg Qual and Med Qual are the mean and median PHRED quality scores for all passing reads, respectively. Means of %, average and median statistics are recalculated with total base count. Mean and median of N50 are calculated using the final N50 output per subline. 5

S2 **Phenotypes of the 4 major clades.** The phenotype for each of the major clades was determined by features defined by the behavior or characteristics of (select) sublines from the clade previously characterized (Methods) [6]. . . . . . . . . . . . . . . . . . . . . . . . . . . . 5

S3 `TreeHarmonizer` **SNV placement statistics relative to each subline.** Clonal, subclonal and private represent the percentages of placed SNVs that were called in the respective subline. Regenotyped SNVs are those who were assigned the "lost variant" genotype by `TreeHarmonizer`. Total placed is respective to each subline. . . . . . . . . . . . . . . . . . . . 7

S4 `TreeHarmonizer` **SV placement statistics respective to each subline.** Clonal, subclonal and private represent the percentages of placed SVs that were called in the respective subline. Total placed is respective to each subline. . . . . . . . . . . . . . . . . . . . . . . . . . . . 8

S5 **DMRs associated with the red clade that overlap putative promoter or putative proximal enhancers of known genes.** A region is said to be a putative promoter for a gene if it is within -350bp to +50bp of the gene's annotated transcriptional start site (Ensembl Release 102) [10, 5]. Of the 17 putative promoters, 9 are also annotated in the EPDnew (v.003) database of experimentally validated promoters [3, 12]. A region is said to be a putative proximal enhancer for a gene if is has enhancer-like signature [1] and is within 2Kbp of the annotated transcriptional start site [4, 9, 8] of the gene. Note that *Gata2* was an outlier in that its putative promoter was noted to be hypermethylated while its expression has a positive log2 fold change (Figure 5d.); however, this may be due to the hypomethylation of a putative promoter of *Gata2* alternative isoforms, which went undetected as its mean methylation difference is not large enough to meet our criteria (Methods). . . . . . . . . . . . 24

| Subline | # Reads | Total Length | Max Length | Mean Length | N50 | Q20(%) | Q30(%) | Avg Qual | Med Qual |
|---|---|---|---|---|---|---|---|---|---|
| C1 | 3,878,605 | 83,059,872,680 | 281,118 | 21,414.9 | 29,251 | 87.94 | 74.02 | 18.44 | 19.891 |
| C3 | 3,847,028 | 86,209,871,989 | 966,796 | 22,409.5 | 32,971 | 80.13 | 64.80 | 17.95 | 20.000 |
| C4 | 1,883,328 | 37,715,867,826 | 307,236 | 20,026.2 | 29,906 | 80.48 | 63.96 | 16.70 | 18.000 |
| C5 | 3,628,129 | 75,421,594,079 | 821,988 | 20,788.0 | 34,327 | 80.43 | 65.20 | 18.43 | 20.000 |
| C6 | 3,902,522 | 84,360,107,492 | 695,762 | 21,616.8 | 32,074 | 82.14 | 66.47 | 18.21 | 20.000 |
| C7 | 4,822,610 | 79,929,281,043 | 854,291 | 16,573.9 | 30,743 | 80.94 | 64.79 | 18.16 | 19.000 |
| C8 | 4,267,557 | 82,984,330,581 | 247,823 | 19,445.4 | 26,286 | 87.31 | 72.82 | 18.22 | 19.251 |
| C9 | 9,876,652 | 61,339,127,538 | 110,146 | 6,210.5 | 11,383 | 81.41 | 65.28 | 16.14 | 17.000 |
| C10 | 3,855,208 | 75,232,321,981 | 584,728 | 19,514.5 | 30,439 | 84.18 | 69.18 | 18.42 | 20.000 |
| C11 | 3,749,668 | 75,507,014,613 | 215,132 | 20,137.0 | 27,720 | 87.15 | 73.12 | 18.15 | 19.535 |
| C12 | 3,927,940 | 81,589,237,368 | 737,990 | 20,771.5 | 34,927 | 82.63 | 66.80 | 18.34 | 20.000 |
| C13 | 10,839,502 | 75,051,614,400 | 107,524 | 6,923.9 | 11,230 | 82.12 | 65.86 | 16.35 | 17.000 |
| C14 | 5,058,904 | 92,251,479,203 | 198,176 | 18,235.5 | 27,970 | 85.93 | 70.91 | 17.72 | 19.000 |
| C15 | 3,522,049 | 80,912,615,895 | 291,845 | 22,973.2 | 35,038 | 81.63 | 66.08 | 17.46 | 19.000 |
| C16 | 2,645,838 | 50,726,621,539 | 280,336 | 19,172.2 | 33,058 | 74.89 | 57.77 | 16.01 | 18.000 |
| C17 | 3,054,146 | 29,144,016,270 | 116,810 | 9,542.4 | 17,309 | 84.13 | 68.60 | 17.04 | 18.000 |
| C18 | 4,790,854 | 44,290,513,825 | 306,231 | 9,244.8 | 18,063 | 73.86 | 56.17 | 15.95 | 17.000 |
| C19 | 3,623,933 | 78,417,582,081 | 466,186 | 21,638.8 | 34,885 | 86.75 | 73.28 | 18.55 | 20.000 |
| C20 | 3,663,675 | 77,163,089,740 | 607,498 | 21,061.7 | 33,248 | 79.99 | 62.59 | 17.30 | 19.000 |
| C21 | 2,575,410 | 53,705,600,160 | 372,912 | 20,853.2 | 35,252 | 83.86 | 68.11 | 18.04 | 20.000 |
| C22 | 2,370,554 | 51,298,610,931 | 403,499 | 21,639.9 | 34,978 | 84.58 | 69.46 | 18.30 | 20.000 |
| C23 | 3,262,926 | 75,206,706,356 | 394,666 | 23,048.9 | 34,899 | 85.16 | 70.51 | 17.86 | 20.000 |
| C24 | 3,833,398 | 73,141,221,559 | 516,441 | 19,080.0 | 32,724 | 82.88 | 66.80 | 18.46 | 20.000 |
| **Mean** | 4,212,193 | 69,767,752,137 | 429,788 | 18,362.0 | 29,073 | 82.95 | 67.50 | 17.77 | 19.240 |
| **Median** | 3,833,398 | 75,421,594,079 | 372,912 | 20,137.0 | 32,074 | 82.63 | 66.80 | 18.04 | 19.535 |
| Spleen Normal | 13,291,753 | 73,498,824,861 | 143,742 | 5,529.7 | 9,975 | 82.80 | 68.58 | 16.51 | 17.000 |
| Parental | 3,720,422 | 85,913,740,338 | 926,409 | 23,092.5 | 35,411 | 83.92 | 68.75 | 18.53 | 20.000 |

Table S1: **Sequencing statistics for the 23 sublines, spleen, and parental B2905 line.** Q20(%) (and Q30(%)) represents the percentage of bases that have a PHRED quality score of 20 (and 30) or over. Avg Qual and Med Qual are the mean and median PHRED quality scores for all passing reads, respectively. Means of %, average and median statistics are recalculated with total base count. Mean and median of N50 are calculated using the final N50 output per subline.

| Feature \ Clade | Green | Orange | Blue | Red |
|---|---|---|---|---|
| Growth rate in C57BL/6 mice | ++ | + | +/- | +++ |
| Response to host immunity (Delay of growth initiation in C57BL/6) | High | Medium | Very high | Low |
| ICB response (anti-CTLA4 vs. IgG) | Medium | High | (growing too slow for study) | Low |
| Cell cycle gene expression | Medium | High | Medium | Low |
| Wnt signaling gene expression | Mixed | Canonical | | Non-canonical |
| **Phenotype** | **Mixed, Less Sensitive to ICB** | **Proliferative, Sensitive to ICB** | **Immunogenic, Susceptible to Host Immunity** | **Invasive, ICB Resistant** |

Table S2: **Phenotypes of the 4 major clades.** The phenotype for each of the major clades was determined by features defined by the behavior or characteristics of (select) sublines from the clade previously characterized (Methods) [6].

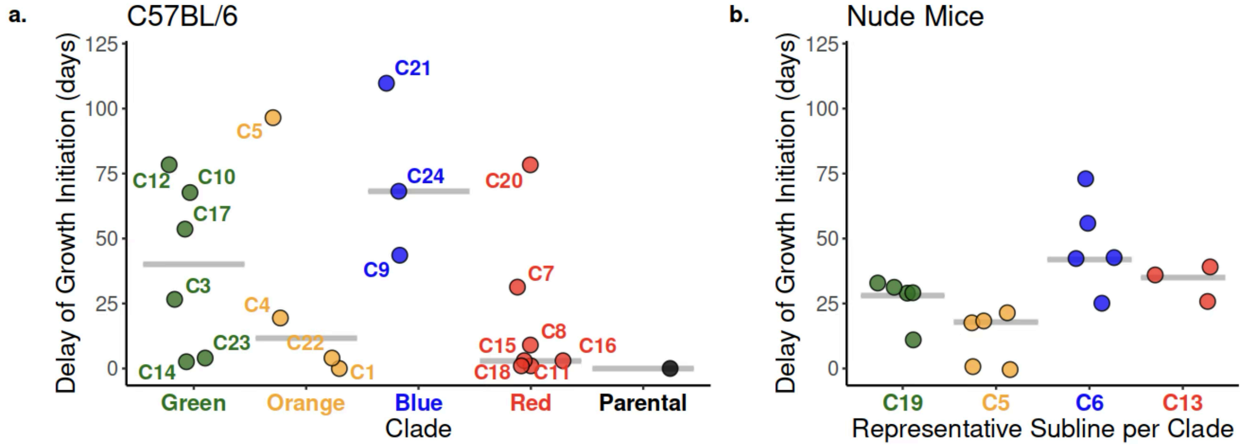


Figure S1: **Delay of growth initiation (DGI) of sublines in C57BL/6 and nude mice.** DGI is defined as the number of days between cell implantation in a mouse and the tumor started continuous growing without regression. **a.** One million cells of each subline were implanted into each of the five C57BL/6 mice (Day 1). The tumor size was measured twice a week. The mean DGI of the sublines in each clade, as well as the median across sublines means within a clade, was shown. No tumor was grown from any mice implanted with cells of C19, C6, or C13. Note that two mice implanted with C5 cells did not grow tumor, and they were not included in the calculation of mean DGI. These results suggested that, on average, sublines in blue clade induced the stronger host immunity than those in other clades. **b.** To investigate the association of the DGI with host immunity, the subline from each clade with the highest DGI following C57BL/6 implantation was tested in immunocompromised host. For each of C19 (green clade), C5 (orange clade), C6 (blue clade), and C13 (red clade), one million cells were implanted into each of the five immunocompromised nude mice (Day 1), and the DGI values measured from each mouse, as well as the median across mouse DGI values for a subline are shown. As expected, we observed the tumor grew in all mice with diminishing DGI as compared to the implantation in C57BL/6 mice. These results indicated that DGI is associated with subline response to host immunity.

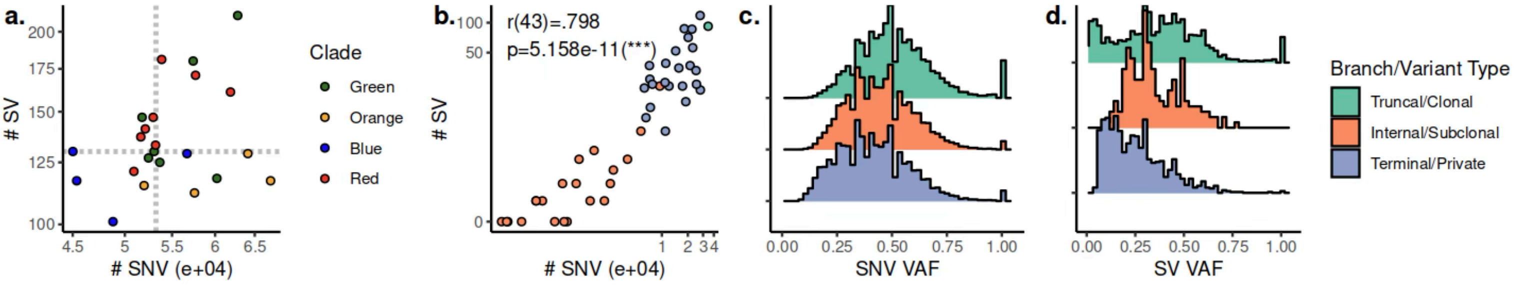


Figure S2: **SNVs and SVs placed by `TreeHarmonizer`. a.** Cumulative number of placed SNVs and SVs per subline, colored by clade. Medians are marked by dashed lines. **b.** The number of SNVs and the number of SVs placed at each branch of the phylogeny strongly correlate with each other. **c.** Variant allele frequency distributions of placed SNVs. **d.** Variant allele frequency distributions of placed SVs.

| Subline | Clonal (%) | Subclonal (%) | Private (%) | Regenotyped | Total Placed |
|---|---|---|---|---|---|
| C1 | 50.92 | 6.91 | 42.17 | 3 | 66,172 |
| C3 | 63.47 | 13.55 | 22.98 | 3 | 52,661 |
| C4 | 57.02 | 7.89 | 35.09 | 8 | 53,905 |
| C5 | 65.56 | 0.57 | 33.86 | 2 | 51,199 |
| C6 | 76.19 | 7.59 | 16.22 | 0 | 44,258 |
| C7 | 67.21 | 19.62 | 13.17 | 2 | 49,692 |
| C8 | 64.48 | 14.16 | 21.36 | 57 | 51,922 |
| C9 | 59.46 | 1.97 | 38.56 | 0 | 55,089 |
| C10 | 65.43 | 3.69 | 30.88 | 1566 | 50,932 |
| C11 | 59.27 | 5.37 | 35.36 | 2 | 57,140 |
| C12 | 66.47 | 5.05 | 28.48 | 19 | 50,743 |
| C13 | 62.08 | 2.52 | 35.41 | 11 | 51,401 |
| C14 | 64.26 | 14.15 | 21.59 | 3 | 50,411 |
| C15 | 65.06 | 22.53 | 12.41 | 4 | 50,482 |
| C16 | 53.68 | 7.60 | 38.72 | 2 | 58,846 |
| C17 | 53.49 | 1.15 | 45.37 | 7 | 55,490 |
| C18 | 62.74 | 22.53 | 14.74 | 1 | 49,929 |
| C19 | 51.04 | 1.06 | 47.90 | 50 | 57,948 |
| C20 | 66.18 | 19.31 | 14.51 | 11 | 50,030 |
| C21 | 69.88 | 6.90 | 23.22 | 1 | 47,280 |
| C22 | 52.74 | 7.26 | 39.99 | 9 | 62,225 |
| C23 | 59.29 | 1.31 | 39.40 | 0 | 56,589 |
| C24 | 75.39 | 7.21 | 17.40 | 2 | 44,618 |
| Mean | 62.23 | 8.69 | 29.08 | 76 | 52,998 |
| Median | 63.47 | 7.21 | 30.88 | 3 | 51,401 |

Table S3: `TreeHarmonizer` **SNV placement statistics relative to each subline.** Clonal, subclonal and private represent the percentages of placed SNVs that were called in the respective subline. Regenotyped SNVs are those who were assigned the "lost variant" genotype by `TreeHarmonizer`. Total placed is respective to each subline.

| Subline | Clonal (%) | Subclonal (%) | Private (%) | Total Placed |
|---|---|---|---|---|
| C1 | 77.69 | 3.85 | 18.46 | 130 |
| C3 | 73.57 | 7.86 | 18.57 | 140 |
| C4 | 82.40 | 4.00 | 13.60 | 125 |
| C5 | 80.47 | 0.00 | 19.53 | 128 |
| C6 | 67.55 | 1.32 | 31.13 | 151 |
| C7 | 76.12 | 13.43 | 10.45 | 134 |
| C8 | 61.08 | 9.58 | 29.34 | 167 |
| C9 | 67.39 | 0.00 | 32.61 | 138 |
| C10 | 66.88 | 1.95 | 31.17 | 154 |
| C11 | 49.05 | 3.81 | 47.14 | 210 |
| C12 | 54.01 | 1.60 | 44.39 | 187 |
| C13 | 39.64 | 0.90 | 59.46 | 222 |
| C14 | 69.50 | 7.80 | 22.70 | 141 |
| C15 | 64.56 | 20.89 | 14.56 | 158 |
| C16 | 49.75 | 6.90 | 43.35 | 203 |
| C17 | 76.23 | 1.64 | 22.13 | 122 |
| C18 | 65.75 | 23.29 | 10.96 | 146 |
| C19 | 34.77 | 0.72 | 64.52 | 279 |
| C20 | 64.15 | 11.32 | 24.53 | 159 |
| C21 | 88.70 | 1.74 | 9.57 | 115 |
| C22 | 69.13 | 3.36 | 27.52 | 149 |
| C23 | 43.97 | 0.86 | 55.17 | 232 |
| C24 | 75.57 | 1.53 | 22.9 | 131 |
| Mean | 65.13 | 5.58 | 29.29 | 161.78 |
| Median | 67.39 | 3.36 | 24.53 | 149 |

Table S4: `TreeHarmonizer` **SV placement statistics respective to each subline.** Clonal, subclonal and private represent the percentages of placed SVs that were called in the respective subline. Total placed is respective to each subline.

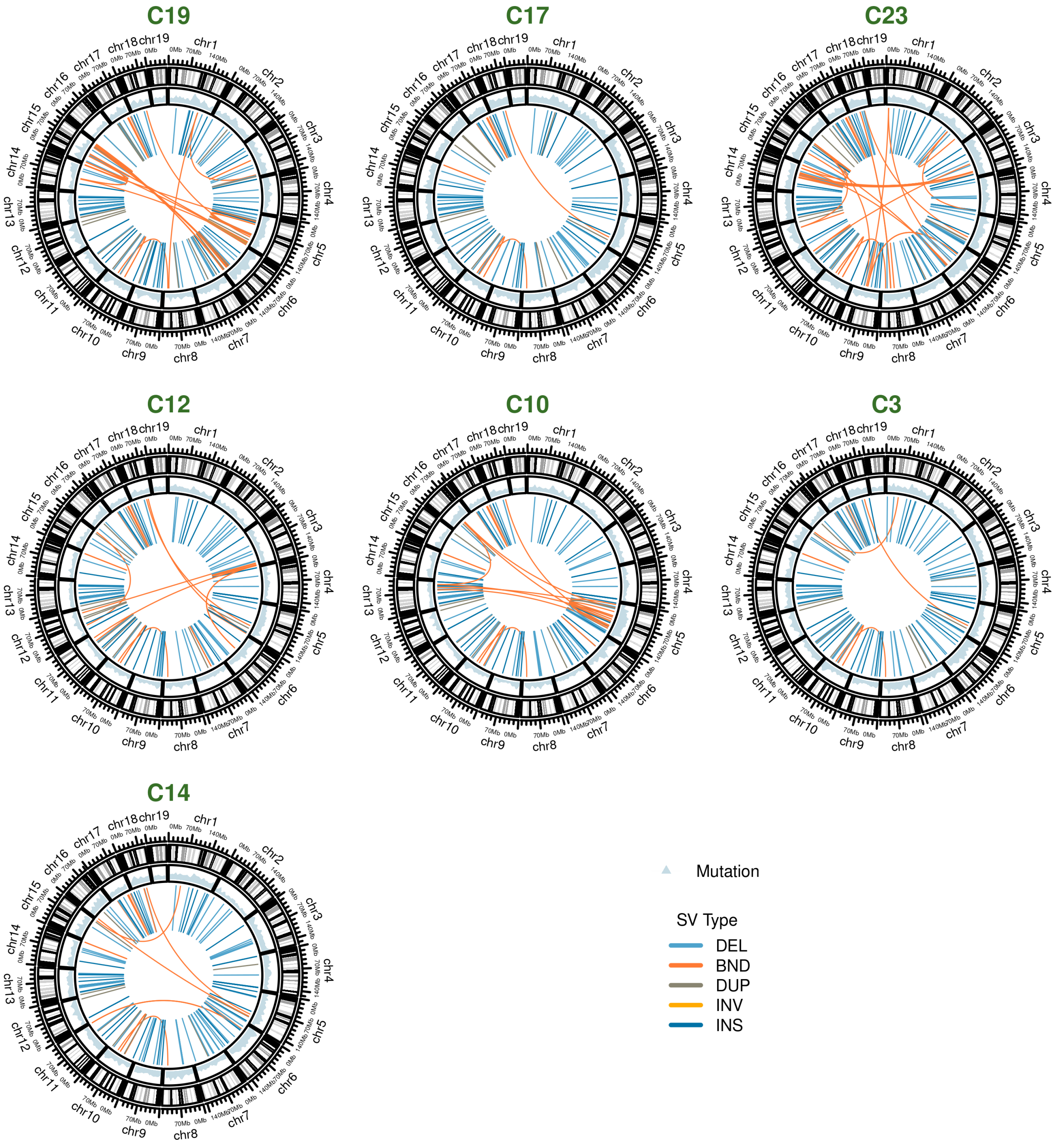


Figure S3: **SNV density and SV events present in individual sublines in the green clade.**

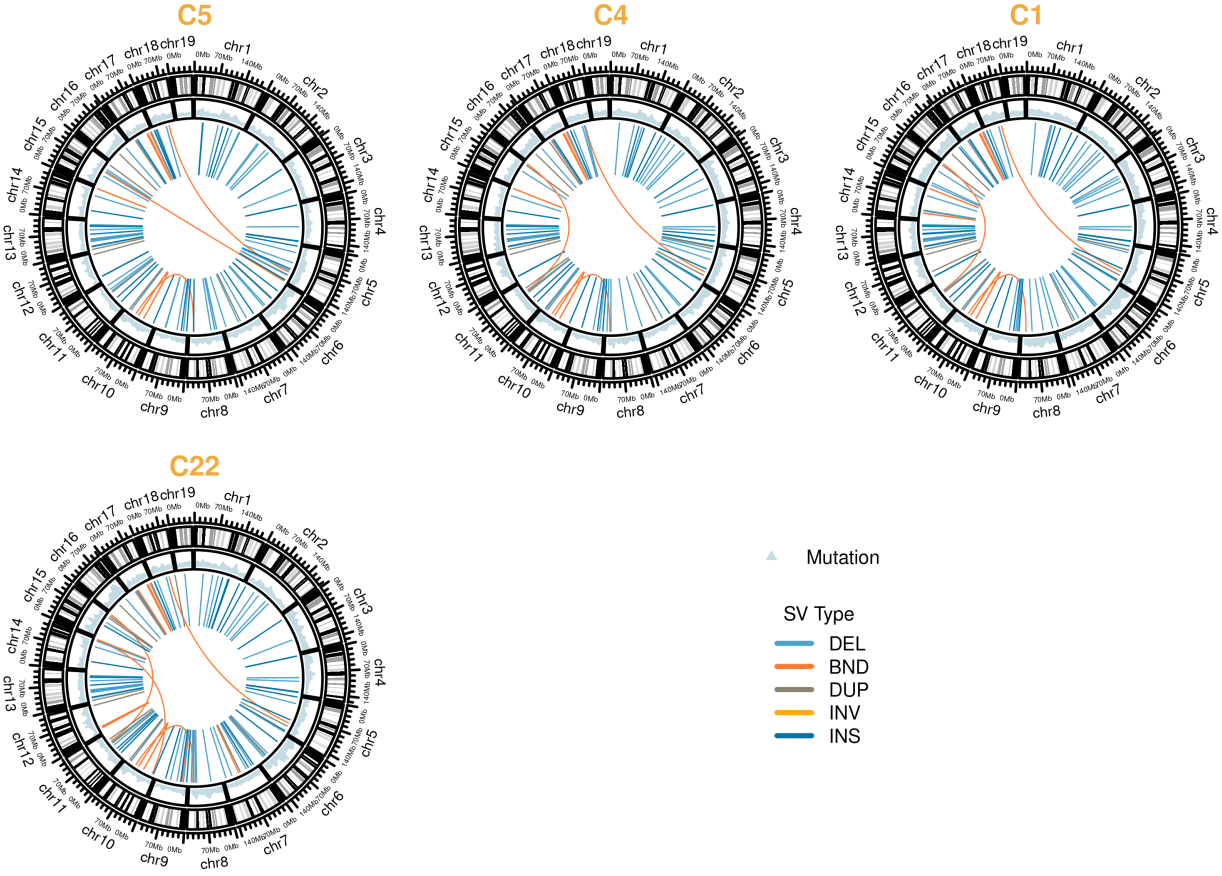


Figure S4: **SNV density and SV events present in individual sublines in the orange clade..**

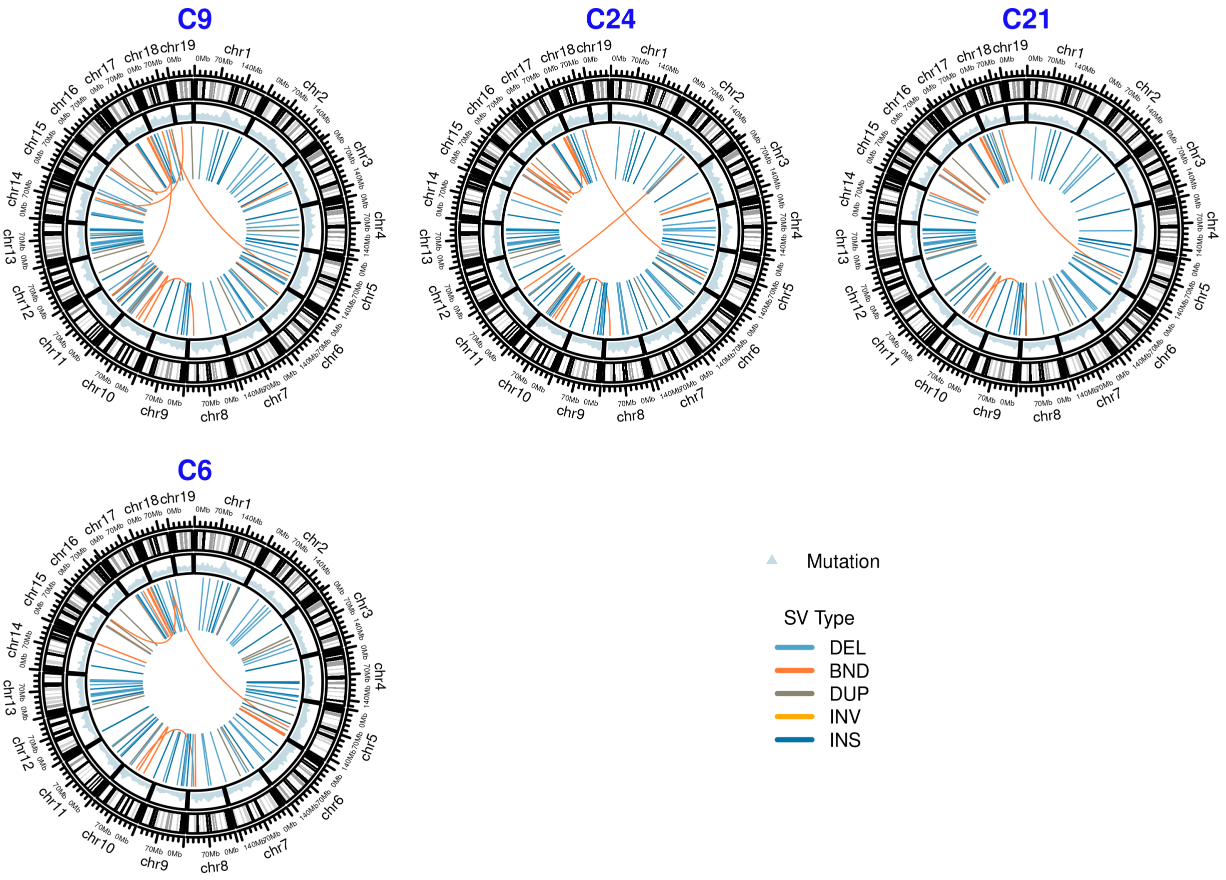


Figure S5: **SNV density and SV events present in individual sublines in the blue clade.**

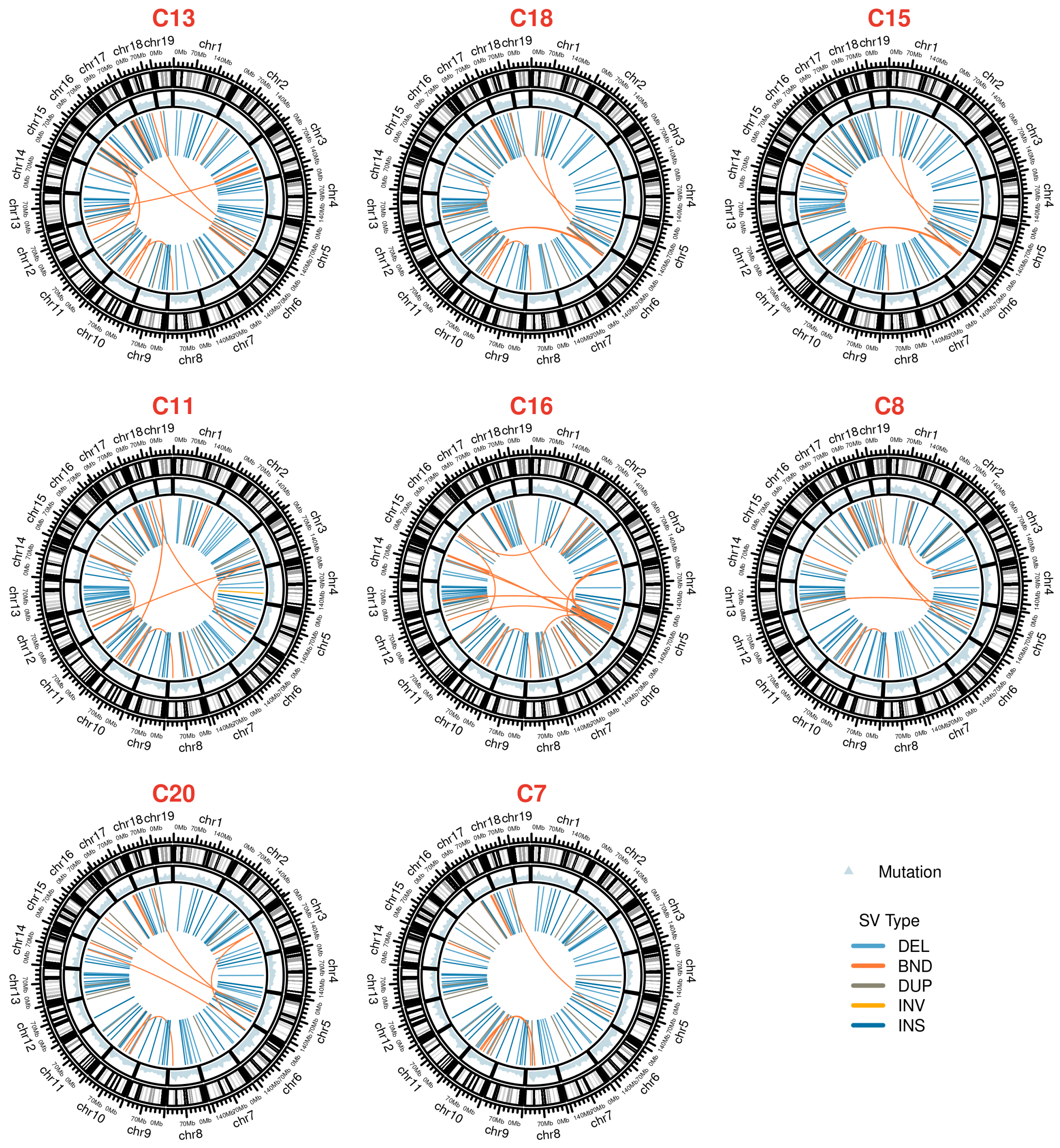


Figure S6: **SNV density and SV events present in individual sublines in the red clade.**

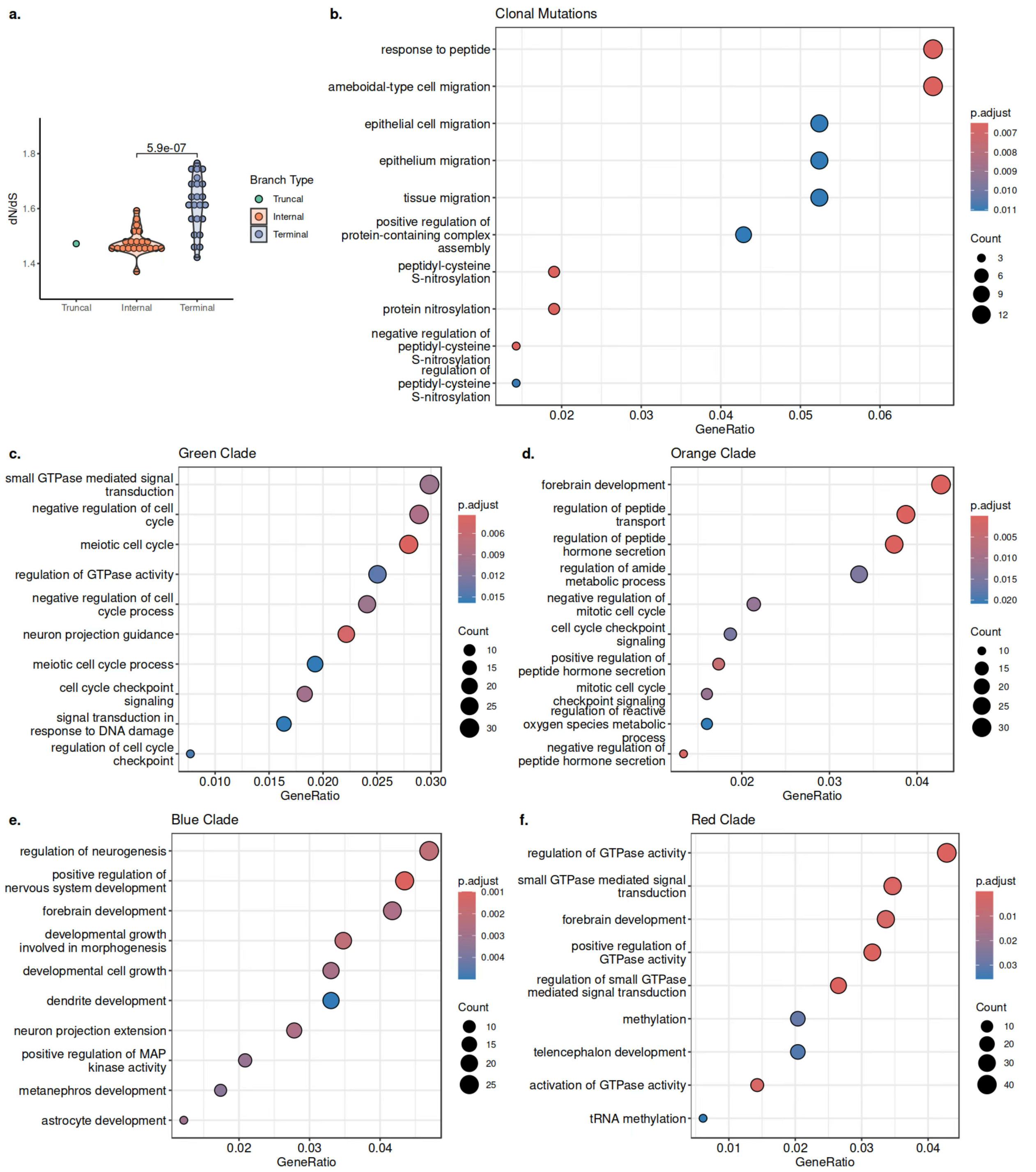


Figure S7: **SNVs along the tumor phylogeny. a.** dN/dS value distributions among truncal, internal, and terminal branches. dN/dS values at internal branches are significantly lower than that at terminal branches by one-tailed *t*-test. **b.** Select enrichment of genes harboring non-synonymous clonal mutations among GO Biological Process term. **c.-f.** Select enrichment of genes uniquely harboring non-synonymous mutations in the green (c.), orange (d.), blue (e.), and red (f.) clade among GO Biological Process terms.

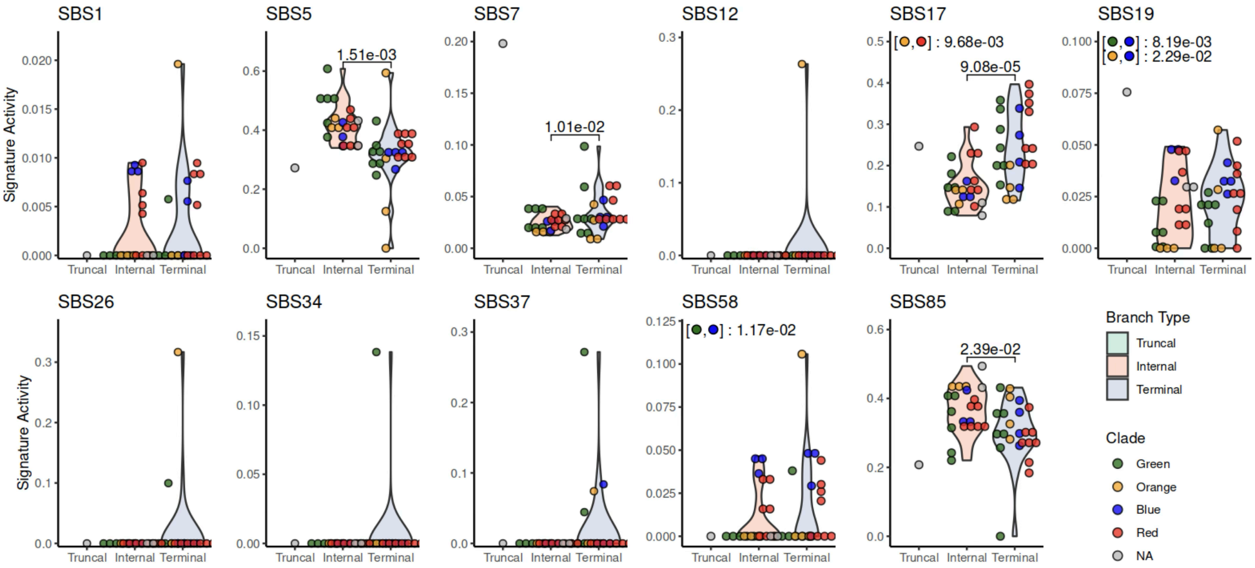


Figure S8: **Mutational signature activities along the tumor phylogeny.** In each panel, a point represents a branch in the phylogeny. A branch can either be categorized into truncal, internal or terminal, or by the color of the clade it is in. ANOVA with post-hoc Tukey HSD (Honestly Significant Difference) Test was performed, and the adjusted $p$-values of statistically significant pairs are reported.

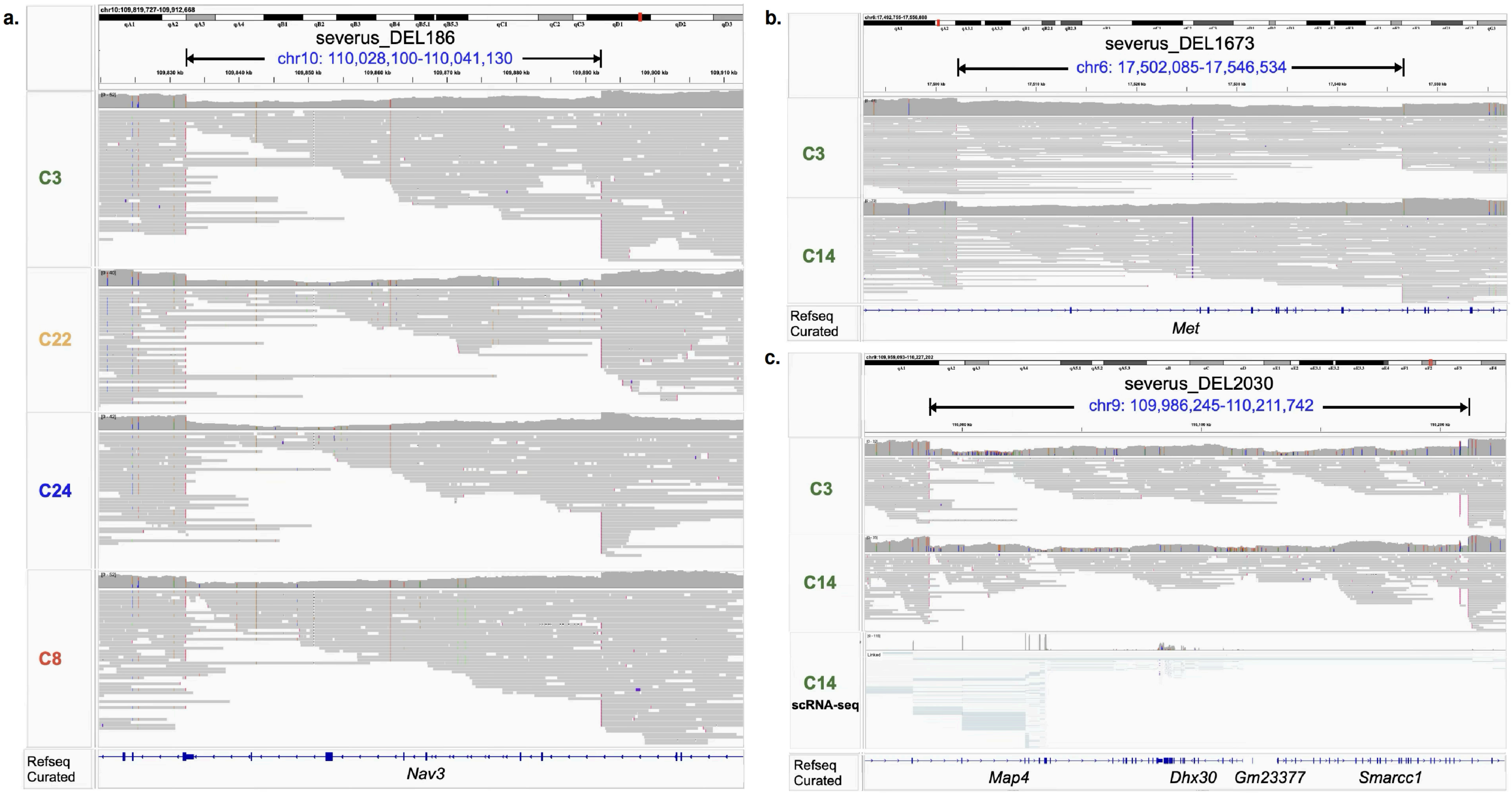


Figure S9: **SV-disrupted genes and gene fusion annotated by `Padfoot`.** **a.** A clonal deletion affects exon 4-9 of *Nav3*. Long-read genomic reads of an representative subline from each major clade are shown. **b.** A subclonal deletion in C3 and C14 in *Met* resulted in the loss of exon 4-11. Long-read genomic reads from sublines C3 and C14 are shown. **c.** A subclonal deletion in C3 and C14 resulted in the fusion of *Map4* and *Smarcc1*. Long-read genomic reads from sublines C3 and C14, as well as scRNA-seq reads from a C14 cell corroborating the fusion event are shown.

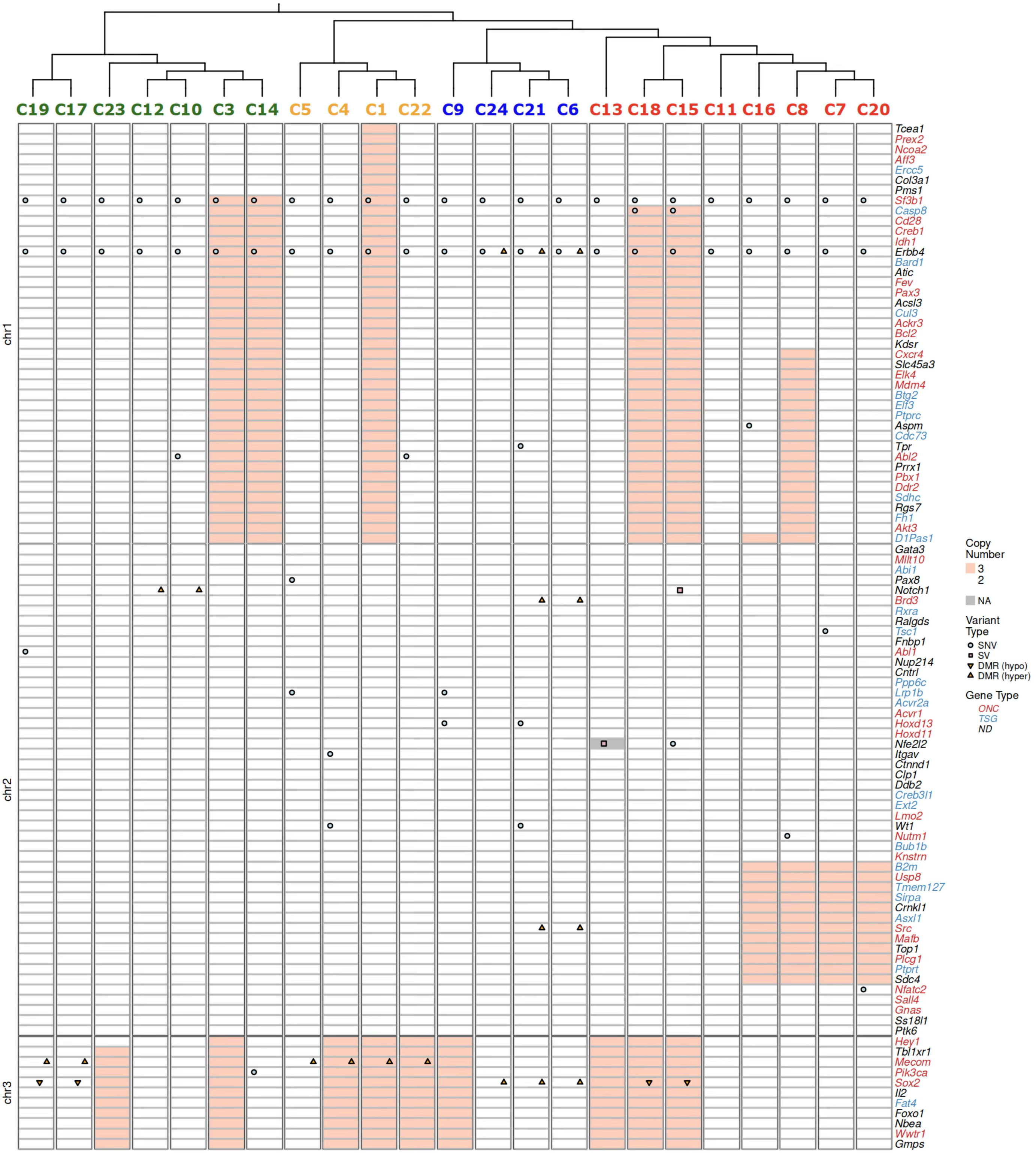


Figure S10: **Mouse homologs of COSMIC genes impacted by (1) non-synonymous SNV, (2) SV breakpoint, (3) CNA, or (4) (hypo/hyper-) DMRs in their (putative) enhancers or promoters in 23 sublines (1/7).** Genes with "NA" copy number in a subline span detected CNA boundaries and therefore cannot be assigned a unique copy number.

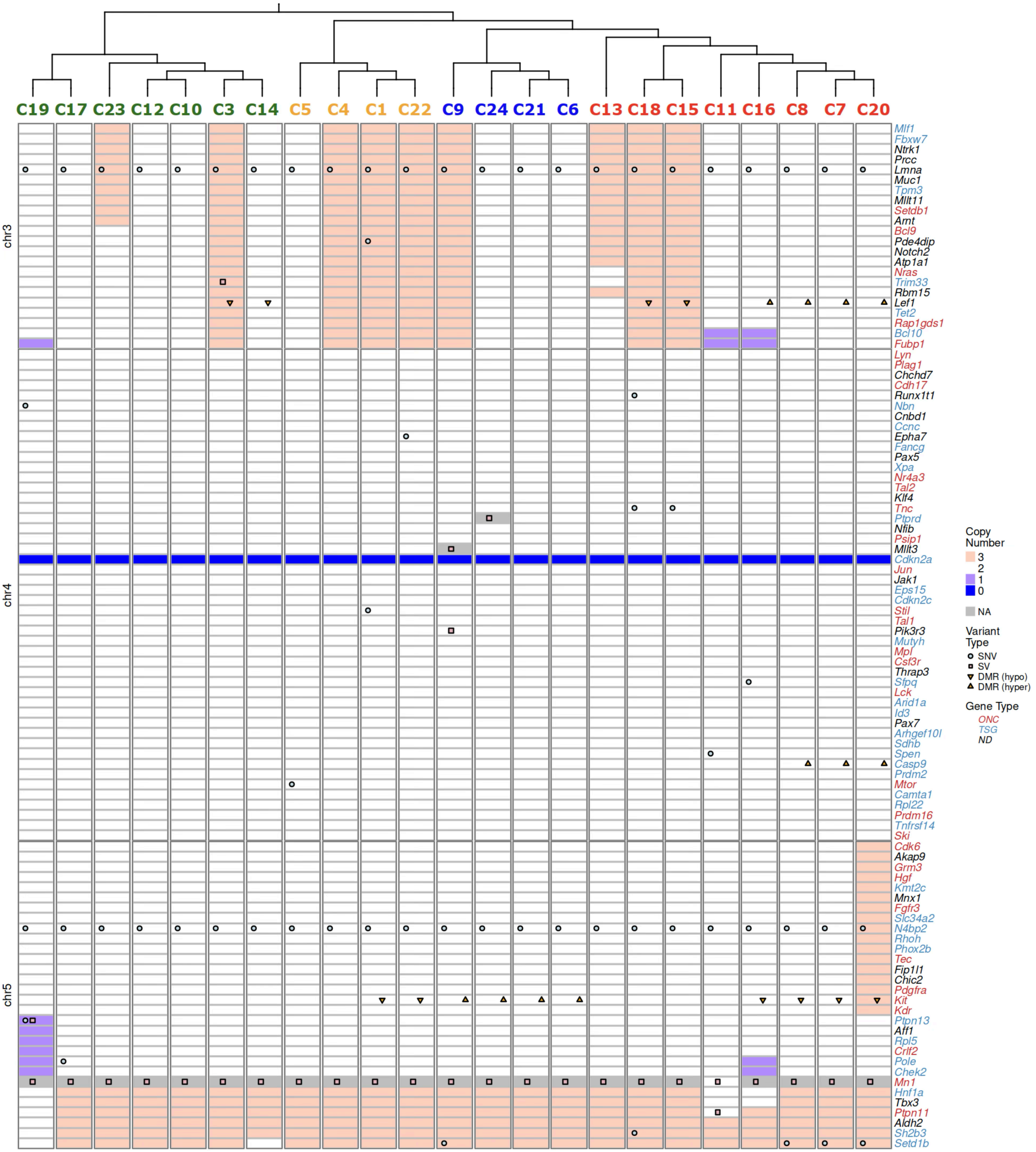


Figure S11: **Mouse homologs of COSMIC genes impacted by (1) non-synonymous SNV, (2) SV breakpoint, (3) CNA, or (4) (hypo/hyper-) DMRs in their (putative) enhancers or promoters in 23 sublines (2/7).** Genes with "NA" copy number in a subline span detected CNA boundaries and therefore cannot be assigned a unique copy number.

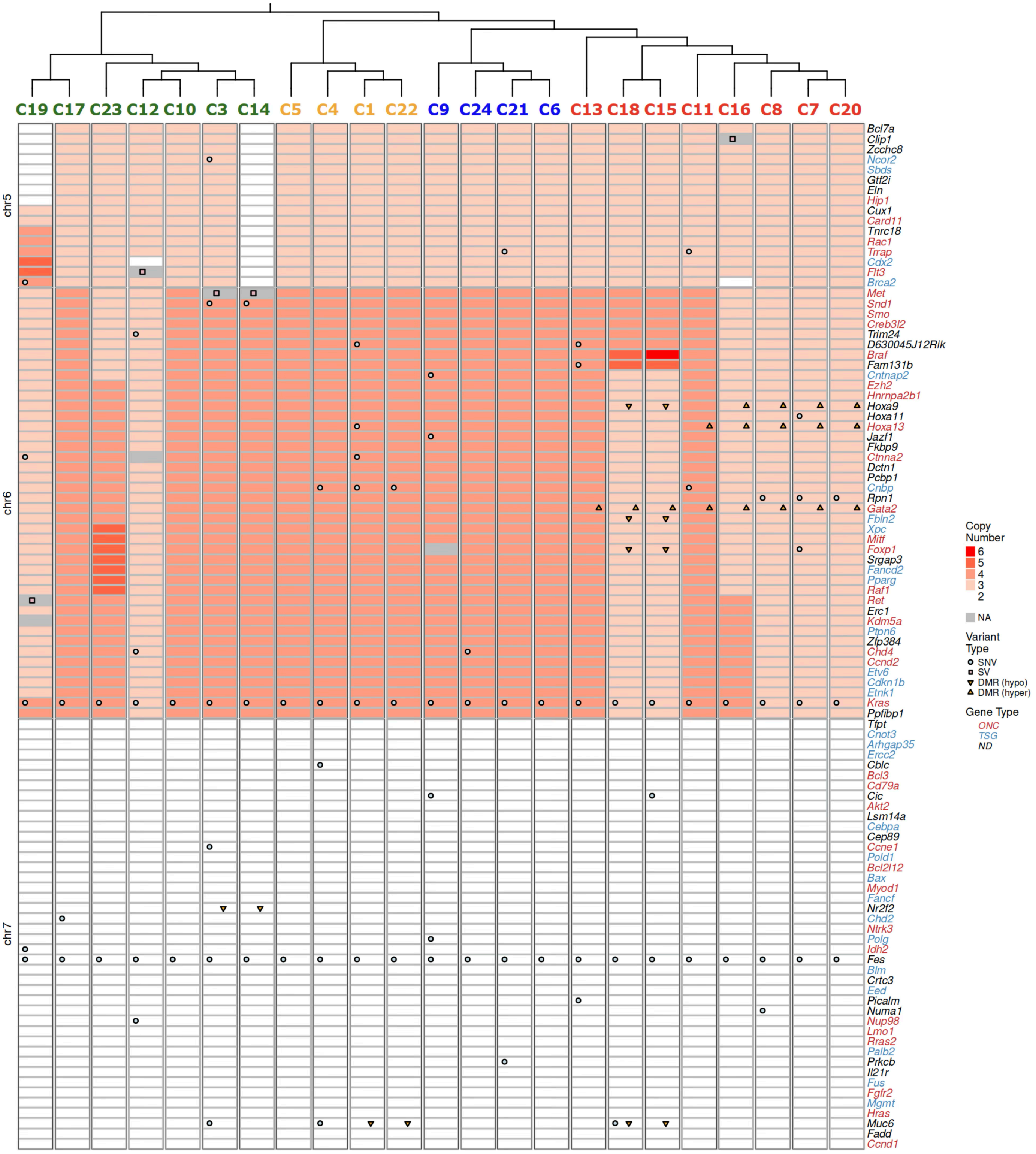


Figure S12: **Mouse homologs of COSMIC genes impacted by (1) non-synonymous SNV, (2) SV breakpoint, (3) CNA, or (4) (hypo/hyper-) DMRs in their (putative) enhancers or promoters in 23 sublines (3/7).** Genes with “NA” copy number in a subline span detected CNA boundaries and therefore cannot be assigned a unique copy number.

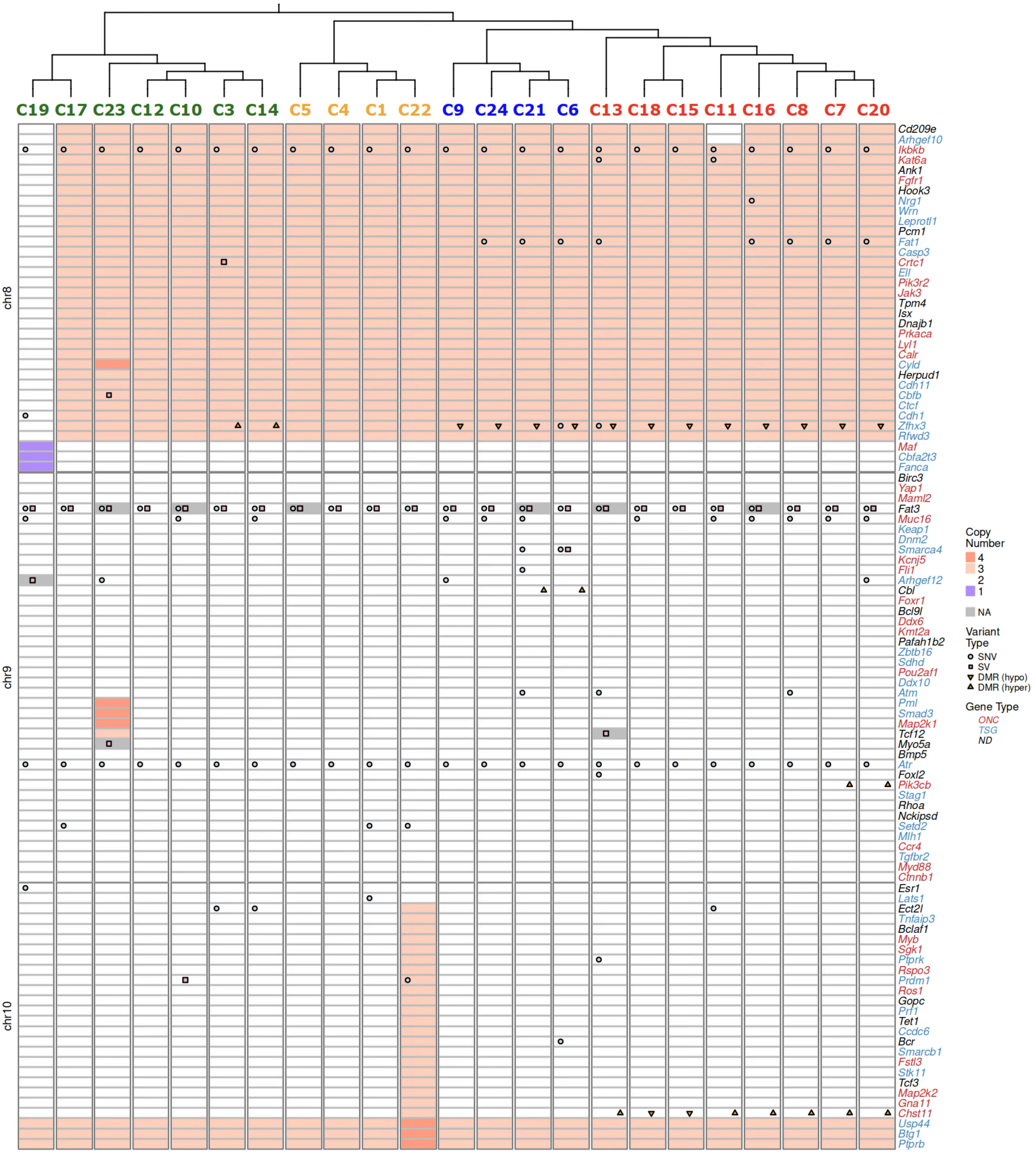


Figure S13: **Mouse homologs of COSMIC genes impacted by (1) non-synonymous SNV, (2) SV breakpoint, (3) CNA, or (4) (hypo/hyper-) DMRs in their (putative) enhancers or promoters in 23 sublines (4/7).** Genes with "NA" copy number in a subline span detected CNA boundaries and therefore cannot be assigned a unique copy number.

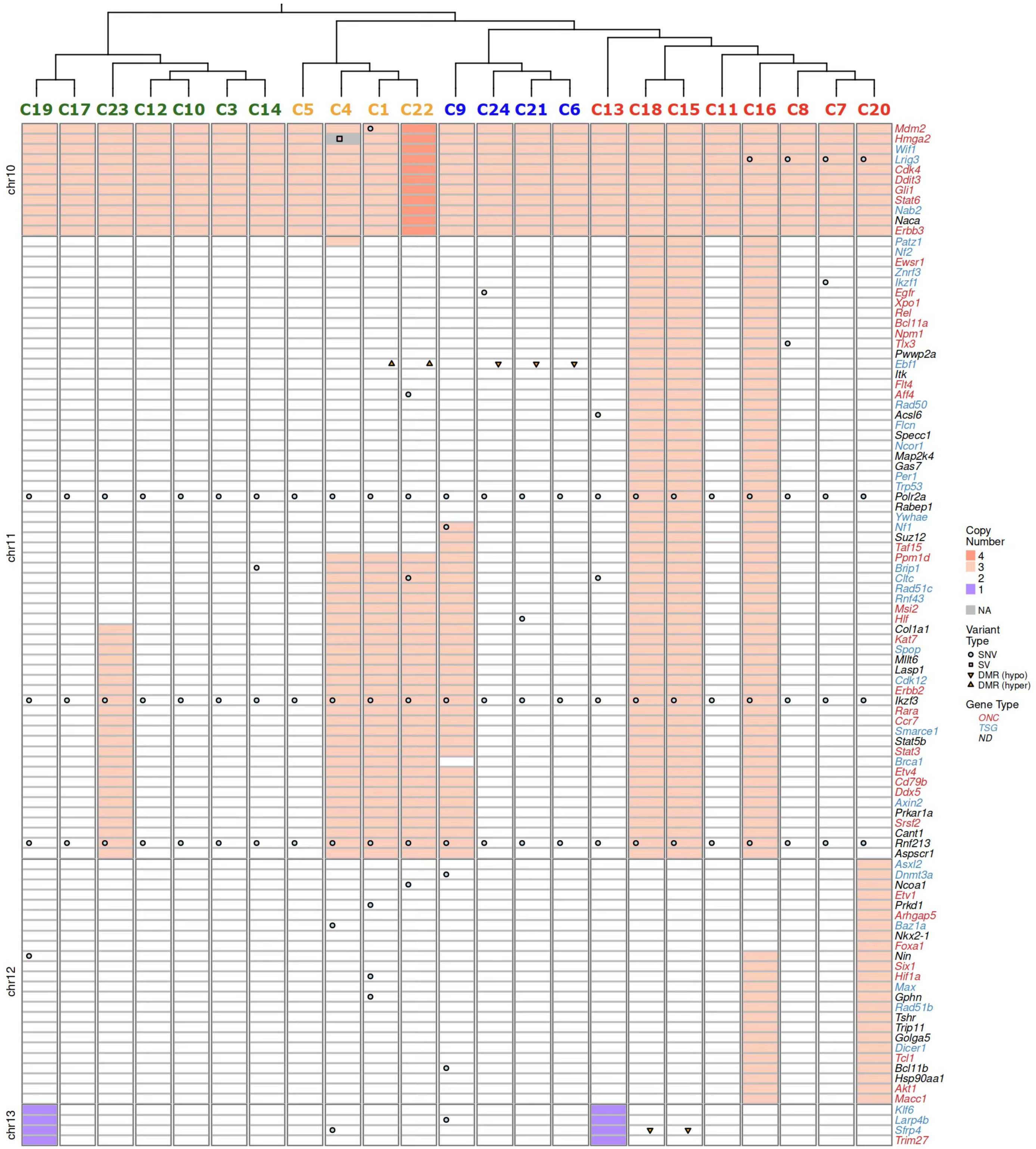


Figure S14: **Mouse homologs of COSMIC genes impacted by (1) non-synonymous SNV, (2) SV breakpoint, (3) CNA, or (4) (hypo/hyper-) DMRs in their (putative) enhancers or promoters in 23 sublines (5/7).** Genes with "NA" copy number in a subline span detected CNA boundaries and therefore cannot be assigned a unique copy number.

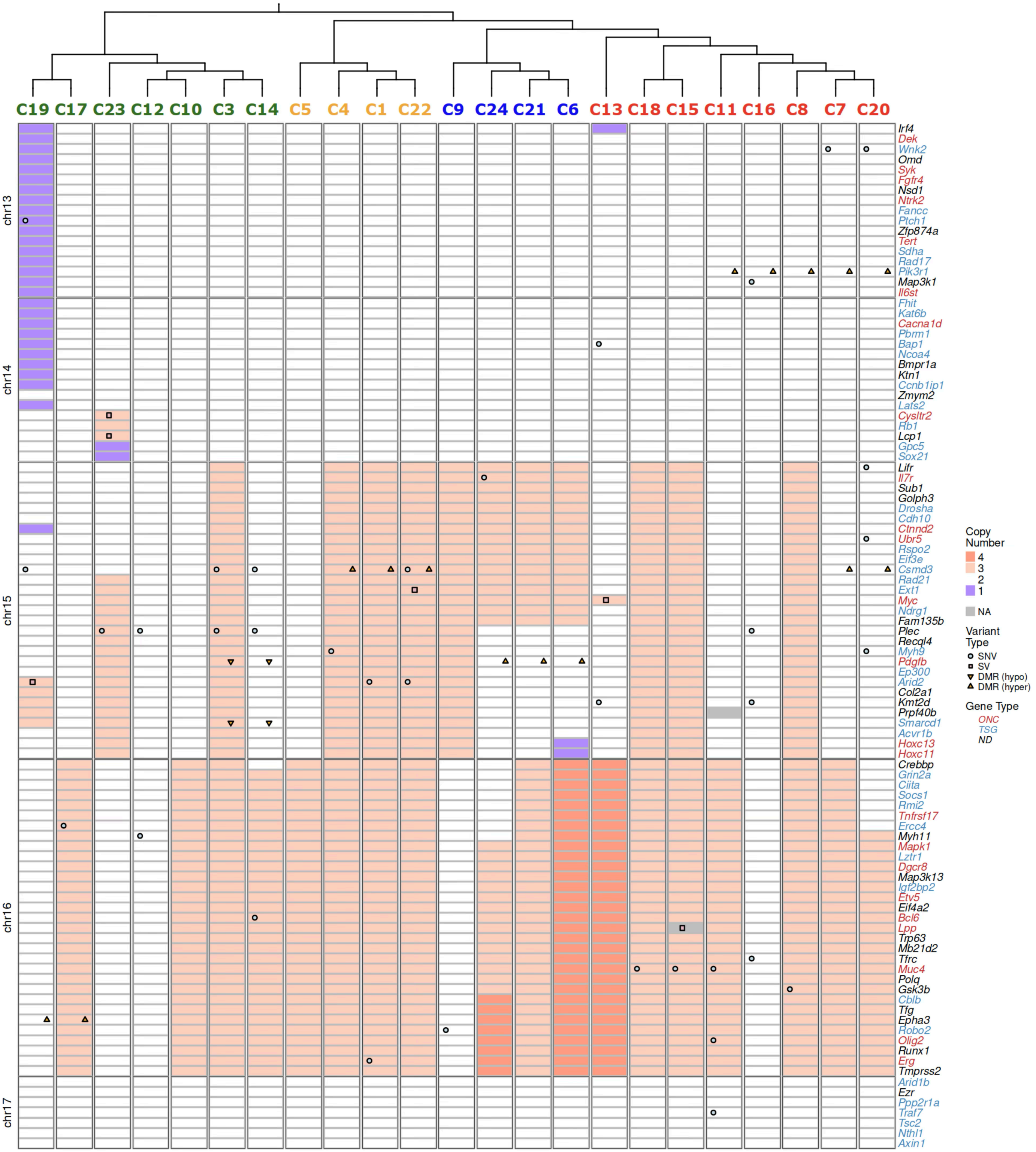


Figure S15: **Mouse homologs of COSMIC genes impacted by (1) non-synonymous SNV, (2) SV breakpoint, (3) CNA, or (4) (hypo/hyper-) DMRs in their (putative) enhancers or promoters in 23 sublines (6/7).** Genes with "NA" copy number in a subline span detected CNA boundaries and therefore cannot be assigned a unique copy number.

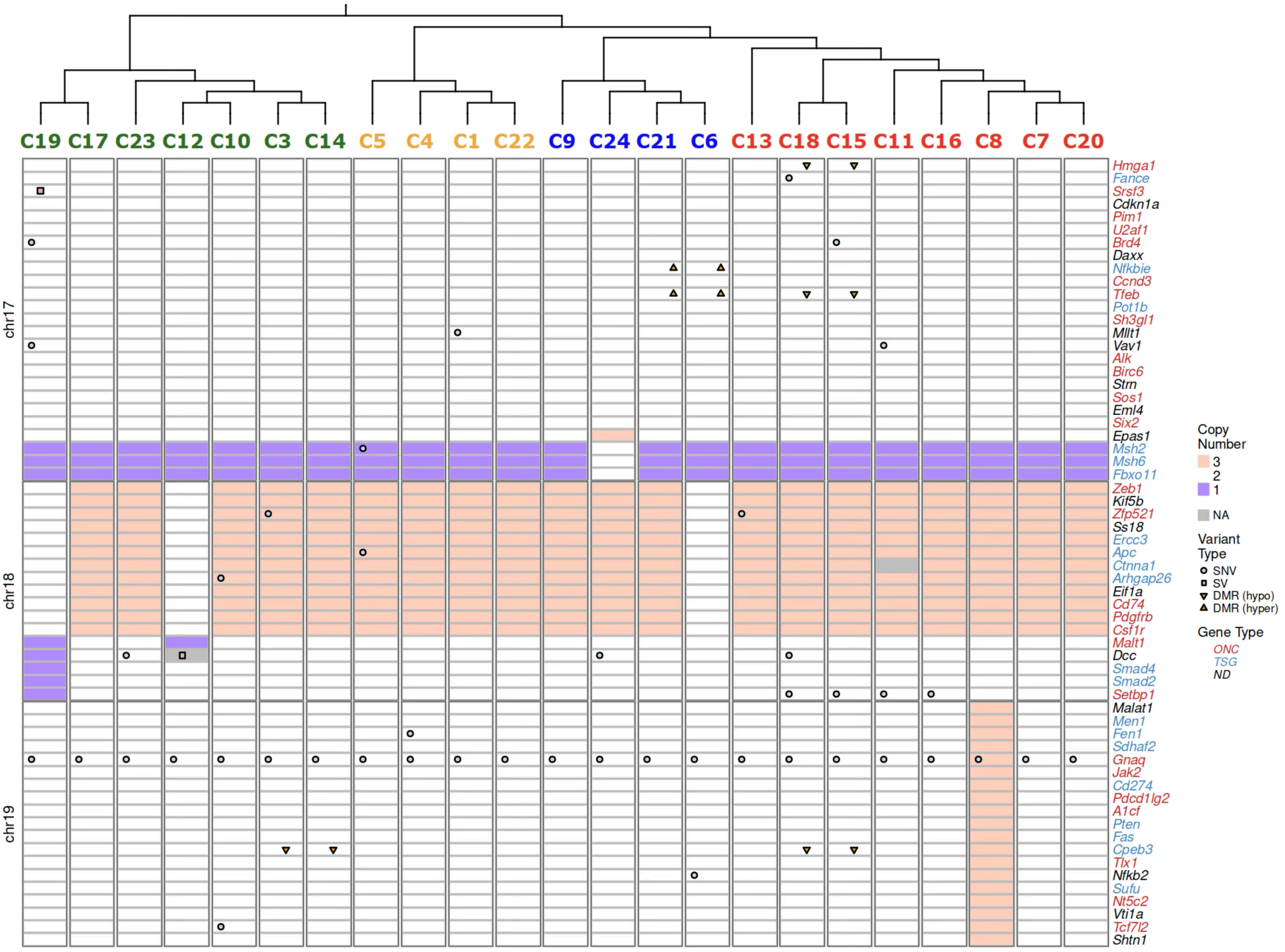

Figure S16: **Mouse homologs of COSMIC genes impacted by (1) non-synonymous SNV, (2) SV breakpoint, (3) CNA, or (4) (hypo/hyper-) DMRs in their (putative) enhancers or promoters in 23 sublines (7/7).** Genes with "NA" copy number in a subline span detected CNA boundaries and therefore cannot be assigned a unique copy number.

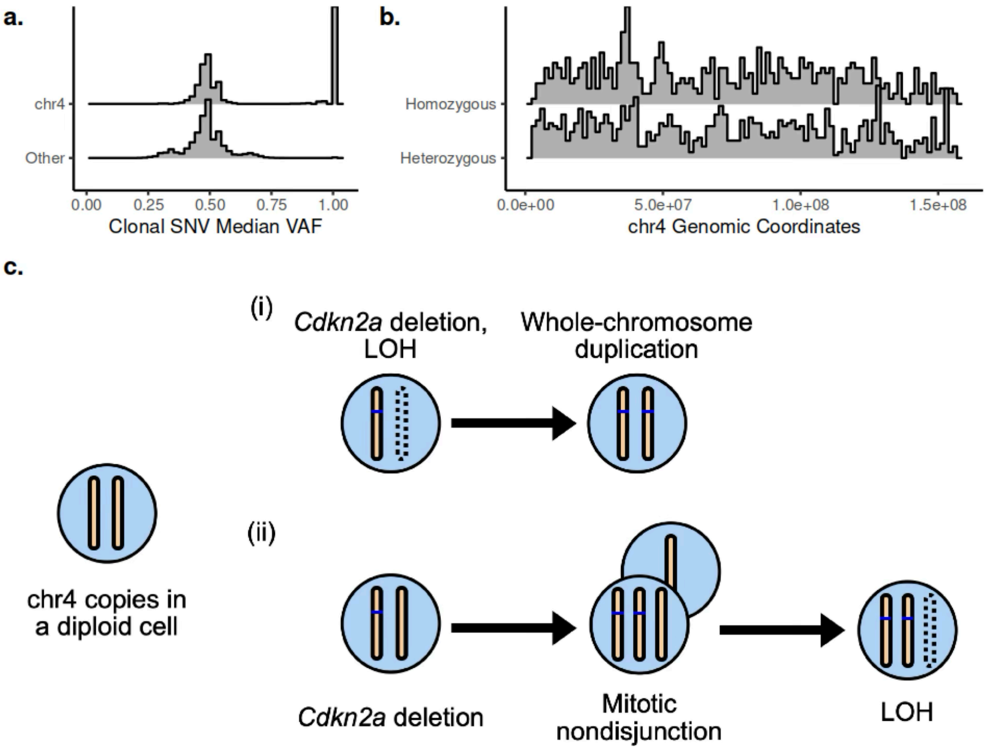


Figure S17: **SNV evidence for the timing clonal CNA events. a.** Median (across sublines) VAF distribution for clonal SNVs on chr4 and other autosomes. **b.** Genomic distributions of homozygous (median VAF = 1) and heterozygous (median VAF < 1) chr4 clonal SNVs. **c.** Two indistinguishable scenarios that may have resulted in the copy-neutral LOH.

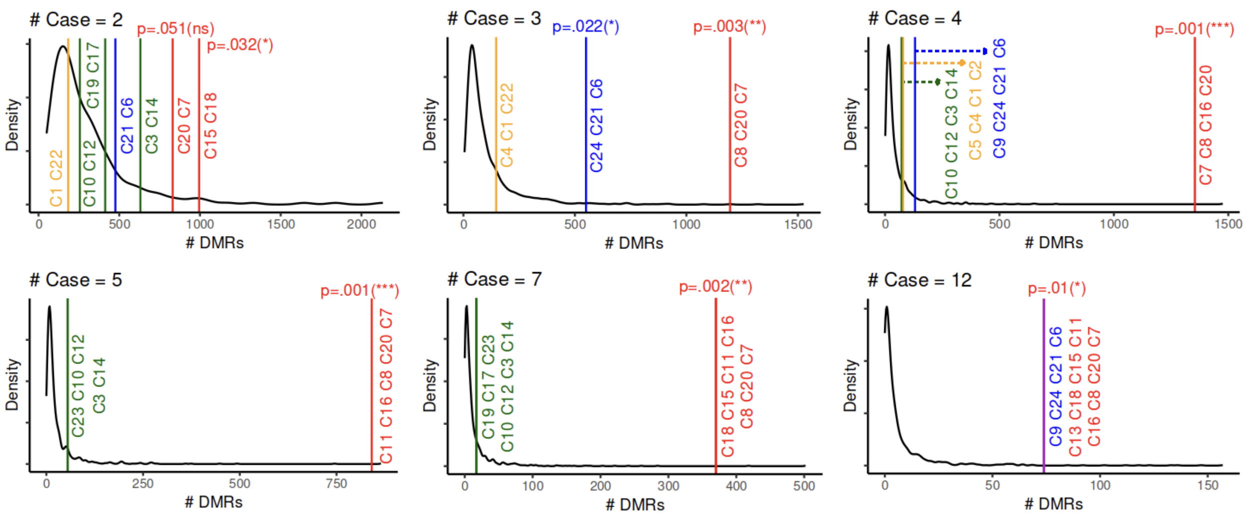


Figure S18: **Permutation tests on the number of DMRs obtained for all subtree sizes of the phylogeny in Figure 2a.** In each panel, "# Case" indicates the number of leaves in the subtree below the branch at which the test was performed. See Figure 5b. for "# Case = 8". Note that due to symmetry, no additional permutation test is done for "# Case = 16" to account for the subtree consisting of the orange, blue, and red in Figure 2a., as the empirical null distribution is identical to that of "# Case = 7" and the number of DMRs obtained is identical to that of the subtree of size 7 colored in green in Figure 2a.

| Chr | Start | End | Direction | Putative Promoter | EPDnew | Proximal Enhancer |
|---|---|---|---|---|---|---|
| chr2 | 156447300 | 156448010 | hyper | *Gm14225* | - | *Epb41l1*, *Gm14225* |
| chr6 | 30736515 | 30739964 | hyper | *Mest* | *Mest* | *Mest*, *Mir335*, *Gm27522*, *Gm27568*, *Gm44296* |
| chr6 | 43264995 | 43267174 | hyper | *Arhgef5*, *Gm44731* | *Arhgef5* | *Arhgef5*, *Gm44731* |
| chr6 | 52203136 | 52204841 | hyper | *Hoxa5* | - | *Hoxa5*, *Hoxa6*, *Hoxaas3*, *Hoxa3*, *Gm28308* |
| chr6 | 73220915 | 73223008 | hyper | *Dnah6*, *Gm40377* | *Dnah6* | *Dnah6*, *Gm40377* |
| chr6 | 88188695 | 88190058 | hyper | - | - | *Gm38708* |
| chr6 | 88193674 | 88194653 | hyper | *Gata2* | - | *Gata2* |
| chr6 | 116628225 | 116629385 | hyper | *Zfp422* | *Zfp422* | *Zfp422*, *Gm44154* |
| chr7 | 23166848 | 23167923 | hyper | *Gm8728* | - | - |
| chr10 | 83140358 | 83141188 | hyper | - | - | *Chst11*, *1700025N21Rik* |
| chr11 | 28582204 | 28583082 | hyper | - | - | *Ccdc85a* |
| chr3 | 79285435 | 79286044 | hypo | - | - | *Rapgef2*, *6430573P05Rik* |
| chr3 | 146220813 | 146221659 | hypo | *Lpar3* | *Lpar3* | *Lpar3* |
| chr4 | 130171758 | 130174329 | hypo | - | - | *Tinagl1* |
| chr4 | 155774166 | 155775696 | hypo | *Vwa1* | *Vwa1* | *Vwa1* |
| chr5 | 14025106 | 14026982 | hypo | *Sema3e* | *Sema3e* | *Sema3e*, *Gm43519* |
| chr6 | 14897121 | 14900597 | hypo | - | - | *Foxp2* |
| chr6 | 117168413 | 117169671 | hypo | *Cxcl12* | *Cxcl12* | *Cxcl12* |
| chr8 | 34524973 | 34526019 | hypo | - | - | *Gm33968* |
| chr9 | 54300511 | 54301797 | hypo | *Gldnos* | - | - |
| chr11 | 65160117 | 65161795 | hypo | *1700086D15Rik* | - | *Arhgap44*, *1700086D15Rik* |
| chr11 | 96279456 | 96280673 | hypo | - | - | *Hoxb8* |
| chr12 | 82586939 | 82588254 | hypo | *Rgs6* | - | - |
| chr15 | 27467528 | 27468632 | hypo | - | - | *Ank* |
| chr17 | 31299560 | 31300475 | hypo | *Slc37a1* | - | - |
| chrX | 52613765 | 52614700 | hypo | *Gpc3* | *Gpc3* | *Gpc3*, *A630012P03Rik* |

Table S5: **DMRs associated with the red clade that overlap putative promoter or putative proximal enhancers of known genes.** A region is said to be a putative promoter for a gene if it is within -350bp to +50bp of the gene's annotated transcriptional start site (Ensembl Release 102) [10, 5]. Of the 17 putative promoters, 9 are also annotated in the EPDnew (v.003) database of experimentally validated promoters [3, 12]. A region is said to be a putative proximal enhancer for a gene if is has enhancer-like signature [1] and is within 2Kbp of the annotated transcriptional start site [4, 9, 8] of the gene. Note that *Gata2* was an outlier in that its putative promoter was noted to be hypermethylated while its expression has a positive log2 fold change (Figure 5d.); however, this may be due to the hypomethylation of a putative promoter of *Gata2* alternative isoforms, which went undetected as its mean methylation difference is not large enough to meet our criteria (Methods).

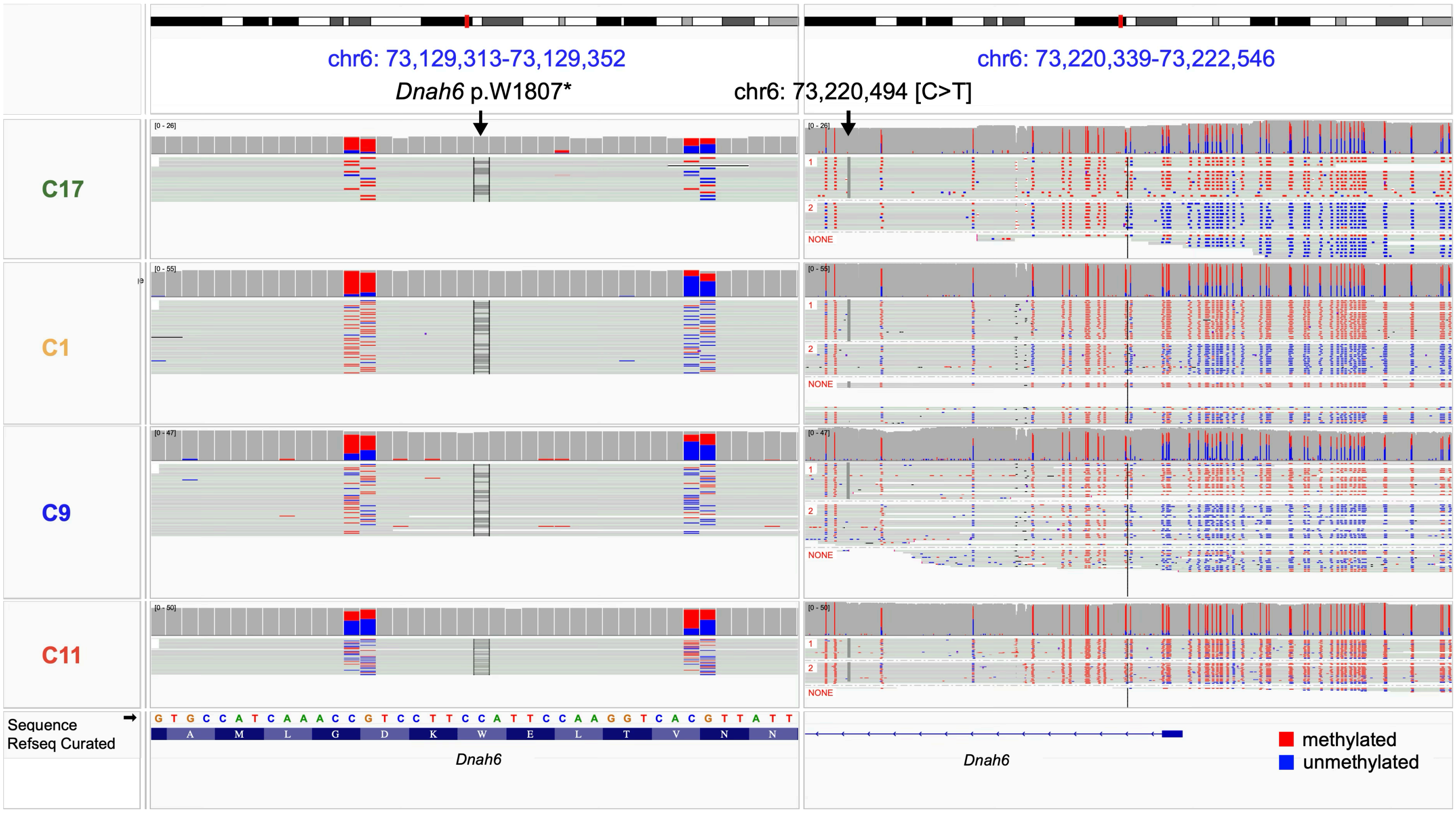


Figure S19: **Associating truncating *Dnah6* mutation with (haplotype-specific) methylation of *Dnah6* promoter in major clades.** One representative subline from each major clade is shown. Clonal truncating mutation *Dnah6* p.W1807* is marked in the left panel displaying unphased reads, and a heterozygous variant chr6: 73,220,494 [C>T] is marked on the right panel with reads phased according to the variant. In C11, the high VAF of *Dnah6* p.W1807*, together with the homozygosity of chr6: 73,220,494 [C>T], not only indicates LOH in C11, but also provides evidence that the two variants exist on the same haplotype. As the phased reads on the right panel are also colored by DNA methylation, we observe that methylation at *Dnah6* promoter is specific to the haplotype harboring chr6: 73,220,494 [C>T], and hence *Dnah6* p.W1807*. We observe haplotype-specific methylation of the *Dnah6* promoter in the green, orange, and blue clade, and hypermethylation in the red clade. Note that while here we observe a few reads from the wild-type copy of *Dnah6* in C11, we see a complete loss of the wild-type copy of *Dnah6* in all other sublines in the red clade, and along with that a complete hypermethylation of the *Dnah6* promoter.

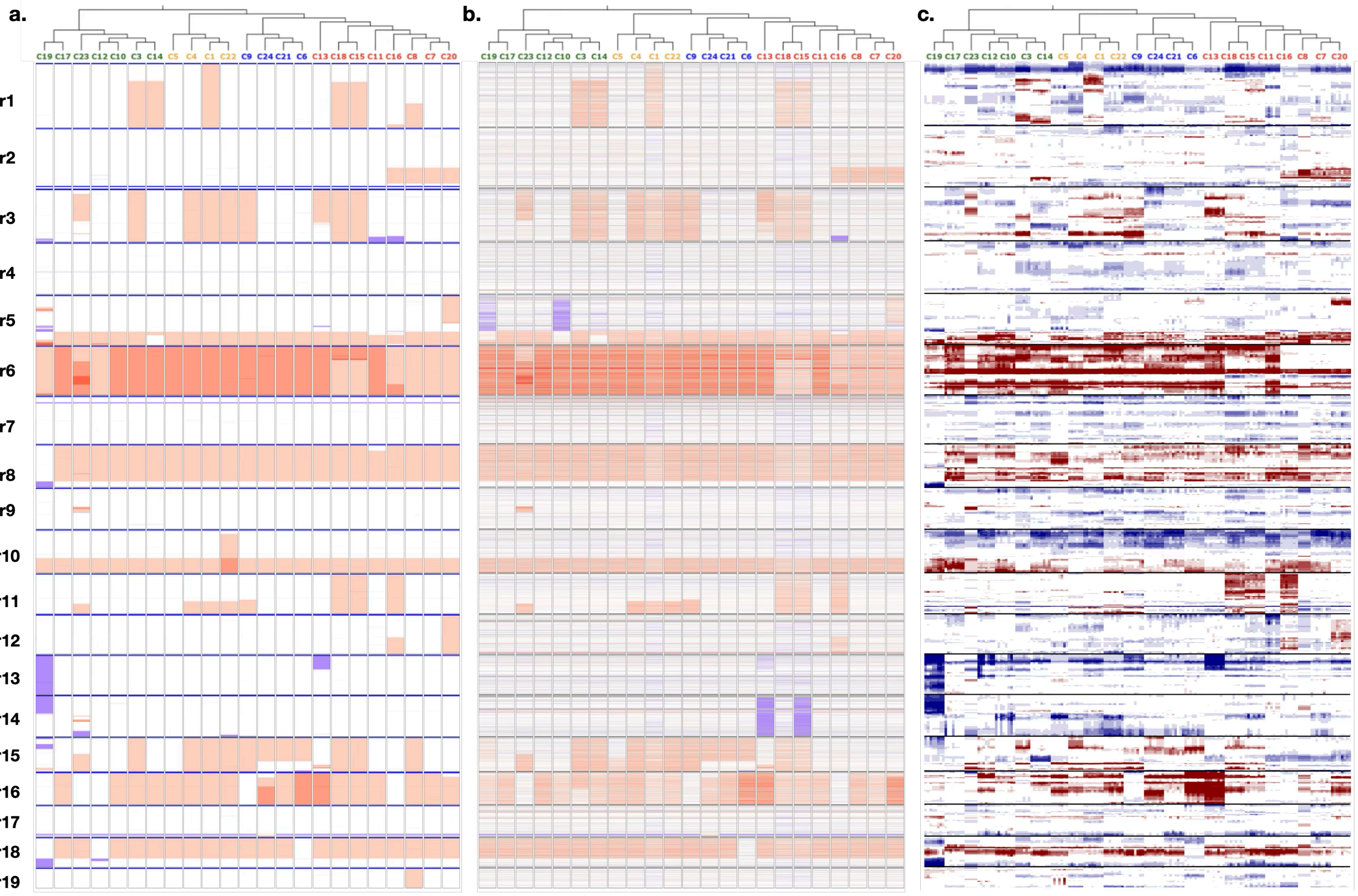


Figure S20: **CNA profiles of 23 sublines obtained using long-read, bWES, and scRNA-seq data.** **a.** Long-read-derived CNA profiles using `Wakhan` [2] (Figure 2e.; Methods) **b.** CNA profiles derived by `CNVkit` [14] from bWES data [11](Methods). **c.** CNA profiles derived by `inferCNV` [15] from full-length scRNA-seq data [7] obtained using the Smart-seq2 protocol [13] (Methods).

[10] F. J. Martin, M. R. Amode, A. Aneja, O. Austine-Orimoloye, A. G. Azov, I. Barnes, A. Becker, R. Bennett, A. Berry, J. Bhai, S. K. Bhurji, A. Bignell, S. Boddu, P. R. Branco Lins, L. Brooks, S. B. Ramaraju, M. Charkhchi, A. Cockburn, L. Da Rin Fiorretto, C. Davidson, K. Dodiya, S. Donaldson, B. El Houdaigui, T. El Naboulsi, R. Fatima, C. G. Giron, T. Genez, G. S. Ghattaoraya, J. G. Martinez, C. Guijarro, M. Hardy, Z. Hollis, T. Hourlier, T. Hunt, M. Kay, V. Kaykala, T. Le, D. Lemos, D. Marques-Coelho, J. C. Marugán, G. A. Merino, L. P. Mirabueno, A. Mushtaq, S. N. Hossain, D. N. Ogeh, M. P. Sakthivel, A. Parker, M. Perry, I. Piližota, I. Prosovetskaia, J. G. Pérez-Silva, A. I. A. Salam, N. Saraiva-Agostinho, H. Schuilenburg, D. Sheppard, S. Sinha, B. Sipos, W. Stark, E. Steed, R. Sukumaran, D. Sumathipala, M.-M. Suner, L. Surapaneni, K. Sutinen, M. Szpak, F. F. Tricomi, D. Urbina-Gómez, A. Veidenberg, T. A. Walsh, B. Walts, E. Wass, N. Willhoft, J. Allen, J. Alvarez-Jarreta, M. Chakiachvili, B. Flint, S. Giorgetti, L. Haggerty, G. R. Ilsley, J. E. Loveland, B. Moore, J. M.

Mudge, J. Tate, D. Thybert, S. J. Trevanion, A. Winterbottom, A. Frankish, S. E. Hunt, M. Ruffier, F. Cunningham, S. Dyer, R. D. Finn, K. L. Howe, P. W. Harrison, A. D. Yates, and P. Flicek. Ensembl 2023. *Nucleic Acids Research*, 51(D1):D933–D941, 11 2022.

[11] F. R. Mehrabadi, E. S. Azer, J. D. Bridgers, E. Pérez-Guijarro, K. L. Marie, H. H. Yang, C. Gruen, C. H. Wu, W. Robinson, H. Liu, C. Kizilkale, M. C. Kelly, C. Smith, S. Chin, J. Ebersole, S. Burkett, A. Buluc, M. P. Lee, E. K. Molloy, T. M. Przytycka, G. Merlino, C.-P. Day, S. Malikić, F. Ergun, and S. C. Sahinalp. A partition function algorithm to evaluate inferred subclonal structures in single-cell sequencing data. In S. Sankararaman, editor, *Research in Computational Molecular Biology*, pages 409–413, Cham, 2025. Springer Nature Switzerland.

[12] P. Meylan, R. Dreos, G. Ambrosini, R. Groux, and P. Bucher. EPD in 2020: enhanced data visualization and extension to ncRNA promoters. *Nucleic Acids Research*, 48(D1):D65–D69, 11 2019.

[13] S. Picelli, O. R. Faridani, Å. K. Björklund, G. Winberg, S. Sagasser, and R. Sandberg. Full-length rna-seq from single cells using smart-seq2. *Nature Protocols*, 9(1):171–181, 2014.

[14] E. Talevich, A. H. Shain, T. Botton, and B. C. Bastian. Cnvkit: Genome-wide copy number detection and visualization from targeted dna sequencing. *PLOS Computational Biology*, 12(4):1–18, 04 2016.

[15] T. Tickle, I. Tirosh, C. Georgescu, M. Brown, and B. Hass. infercnv of the trinity ctat project.
